# Phylogenomics resolves key relationships in *Rumex* and uncovers a dynamic history of independently evolving sex chromosomes

**DOI:** 10.1101/2023.12.13.571571

**Authors:** Mark S. Hibbins, Joanna L. Rifkin, Baharul I. Choudhury, Olena Voznesenka, Bianca Sacchi, Meng Yuan, Yunchen Gong, Spencer C. H. Barrett, Stephen I. Wright

## Abstract

Sex chromosomes have evolved independently many times across eukaryotes. Despite a considerable body of literature on the evolution of sex chromosomes, the causes and consequences of variation in the formation, degeneration, and turnover of sex chromosomes remain poorly understood. Comparative approaches in groups with sexual system variation can be valuable for understanding these questions. Plants are well-suited to such comparative studies, with many lineages containing relatively recent origins of dioecy and sex chromosomes as well as hermaphroditic close relatives. *Rumex* is a diverse genus of flowering plants harboring significant sexual system variation, including hermaphroditic and dioecious clades with XY sex chromosomes. Previous disagreement in the phylogenetic relationships among key species have rendered the history of sex chromosome evolution uncertain. Resolving this history is important to the development of *Rumex* as a system for the comparative study of sex chromosome evolution. Here, we leverage new transcriptome assemblies from 11 species representing the major clades in the genus, along with a whole-genome assembly generated for a pivotal hermaphroditic species, to further resolve the phylogeny and history of sex chromosome evolution in *Rumex*. Using phylogenomic approaches, we find evidence for two independent origins of sex chromosomes and introgression from unsampled taxa in the genus. Comparative genomics reveals massive chromosomal rearrangements in a dioecious species, with evidence for a complex origin of the sex chromosomes through multiple chromosomal fusions. However, we see no evidence of elevated rates of fusion on the sex chromosome in comparison with autosomes, providing no support for an adaptive hypothesis for the sex chromosome expansion. Overall, our results highlight the dynamic nature of sex chromosome systems in *Rumex* and illustrate the utility of the genus as a model for the comparative study of sex chromosome evolution.

## Introduction

Despite the near-universality of sexual reproduction in eukaryotes, the mechanisms that determine an organism’s sex vary greatly (Bull 1983, Bachtrog et al. 2014). In organisms with separate male and female individuals, sex is often genetically determined by the inheritance of sex chromosomes. Classic models propose that sex chromosomes arise from a pair of ancestral autosomes as a resolution to sexually antagonistic selection (Charlesworth & Charlesworth 1980, Rice 1987); suppressed recombination linking antagonistic alleles to the sex in which they are beneficial results in differentiation of the sex chromosomes and often degeneration of the Y or W chromosome over time (Charlesworth 1991, Charlesworth & Charlesworth 2000, Bachtrog 2013). However, these models in isolation are unable to explain why there is so much variation in the age, degree of degeneration, and rate of turnover of sex chromosomes across eukaryotes. Some groups—such as mammals (Graves & Watson 1991, Hughes & Page 2015) and birds (Fridolfsson et al. 1998, Handley et al. 2004) —have mostly ancient, highly conserved, and highly differentiated sex chromosomes. Others, such as amphibians, fishes, and flowering plants, have much younger sex chromosomes and exhibit rapid sex chromosome turnover, with polymorphisms among closely related species or even within species (Ming et al. 2011, Jeffries et al. 2018, Taher et al. 2021).

Recent work has begun to address the factors that might promote or inhibit the formation of new sex chromosomes over time. For example, autosomal pairs that contain large regions of suppressed recombination, or harbor sexually antagonistic variation, may be “primed” for becoming sex chromosomes (Bergero et al. 2019, Rifkin et al. 2021, Guo et al. 2022). Additionally, sex chromosomes in earlier stages of differentiation may be more prone to turning over, as the fitness cost of reverting to an autosomal state may not be as high (Bull and Charnov 1977, Pokorná & Kratochvíl 2009, Vicoso 2019, Lenormand et al. 2020, Lenormand & Roze 2022). Testing these ideas empirically requires comparative approaches, in which the ancestral autosomal homologs of sex chromosomes can be identified in closely related species. The first step in such approaches is resolving the history of sex chromosome evolution within the target group, including the number of independent origins and their relative timescales, in the context of a robust phylogeny. It also necessitates the study of relatively young sex chromosomes in which intact autosomal homologs can be identified. This requirement makes it challenging to apply this approach to lineages with ancient sex chromosome systems like those in mammals.

The distinct genetic architecture of sex chromosomes may favor the evolution of large-scale chromosomal rearrangements, contributing significantly to the evolution of karyotypic differences among species (White 1940, Charlesworth et al. 1987, Connallon et al. 2018). Certain kinds of rearrangements, such as inversions and sex chromosome-autosome fusions, are thought to enable rapid linkage of sexually antagonistic variation to the sex-determining region by extending the region of recombination suppression (Charlesworth & Charlesworth 1980, Rice 1987, Charlesworth et al. 2005). While the importance of structural rearrangements for the formation of new evolutionary strata (Lahn and Page 1999, Handley et al. 2004, Bergero et al. 2007) and neo-sex chromosomes (Kitano et al. 2009, Pala et al. 2012, Castillo et al. 2014, Bracewell et al. 2017) is increasingly well understood, their contributions to macroevolutionary patterns of sex chromosome variation remain understudied. Such comparative approaches can, for example, test whether sex chromosome fusions occur at a higher rate than autosomal fusions, as is expected if fusions with sex chromosomes are favoured as recombination modifiers due to sexually antagonistic selection (Charlesworth & Charlesworth 1980, Anderson et al. 2020). Comparative genomic approaches in a phylogenetic context can be valuable for understanding such processes of karyotype evolution.

Flowering plants (angiosperms) are well-suited for investigations of sex chromosome evolution and turnover. While most species are hermaphroditic, multiple families have recently evolved dioecy with chromosomal sex determination (Charlesworth 2013, Tree of Sex Consortium 2014). One such group is the docks and sorrels (*Rumex*), a globally distributed genus with approximately 200 described species (Grant et al. 2022). *Rumex* harbors significant sexual system diversity, ranging from hermaphrodites with either perfect or unisexual flowers (monoecy) flowers to dioecy (Löve and Kapoor 1967, Navajas-Pérez et al. 2005). There is also considerable variation in the genetic systems, including species with XY and XYY sex chromosomes and within-species sex chromosome polymorphism (e.g. *R. hastatulus* — Smith 1964, Ming et al. 2011, Hough et al. 2014, Beaudry et al. 2020, Rifkin et al. 2021, Beaudry et al. 2022). Populations of *R. hastatulus* from eastern N. America, in addition to multiple other species in the genus, have evolved an XYY sex chromosome system from an ancestral XY configuration, which occurs in populations from western N. America. This can arise via an X-autosome fusion, as is known to have occurred in *R. hastatulus* (Smith 1964, Grabowska-Joachimiak et al. 2015, Kasjaniuk et al. 2019), or via a Y-chromosome fission. Hermaphroditism is the ancestral state in *Rumex*, making it an excellent system for studying the evolution of sex chromosomes from their ancestral autosomal homologs. Previous work on *R. hastatulus* uncovered widespread recombination suppression on all chromosomes, including in the autosomal homolog of the neo-X chromosome found in eastern populations (Rifkin et al. 2021, Rifkin et al. 2022), as well as a role for sex chromosome differences in shaping barriers to contemporary hybridization (Beaudry et al. 2022). However, sex chromosome evolution across the rest of the genus remains poorly understood.

Previous studies constructed phylogenies of *Rumex* using nuclear and chloroplast markers (Navajas-Pérez et al. 2005, Grant et al. 2022, Koenemann et al. 2023). These studies agree that there are two primary clades with sex chromosomes, an XY clade and an XYY clade, but they disagree on the placement of *R. bucephalophorus*, a hermaphroditic/ gynomonoecious (plants with both hermaphroditic and female flowers) species (see Talavera et al. 2011) that lacks sex chromosomes, relative to these clades (Figure 1). The ITS and chloroplast trees of Navajas-Pérez et al. (2005) place the XY and XYY clades as sister, with *R. bucephalophorus* more distantly related (Figure 1A), whereas the other two chloroplast studies place *R. bucephalophorus* as sister to the XY clade, and more distantly related to the XYY clade (Figure 1B, C). Significantly, these two phylogenetic hypotheses have different implications for the sequence of events in the evolution of sex chromosomes in the genus. The former suggests the possibility of a single origin of XY sex chromosomes (Figure 1A), whereas the latter requires two independent changes: either two origins of XY sex chromosomes (Figure 1B)—which also has support from preliminary transcriptome-based identification of sex-linked genes (Crowson et al. 2017)—or a single origin followed by a loss of sex chromosomes in *R. bucephalophorus* (Figure 1C). Phylogenies constructed from small numbers of genetic markers can be vulnerable to both technical errors and biological sources of uncertainty such as incomplete lineage sorting and introgression (Maddison 1997, Degnan and Rosenberg 2009). Resolving the history of sex chromosome evolution in *Rumex* therefore requires the analysis of genome-scale datasets with modern coalescent approaches.

**Figure 1:**
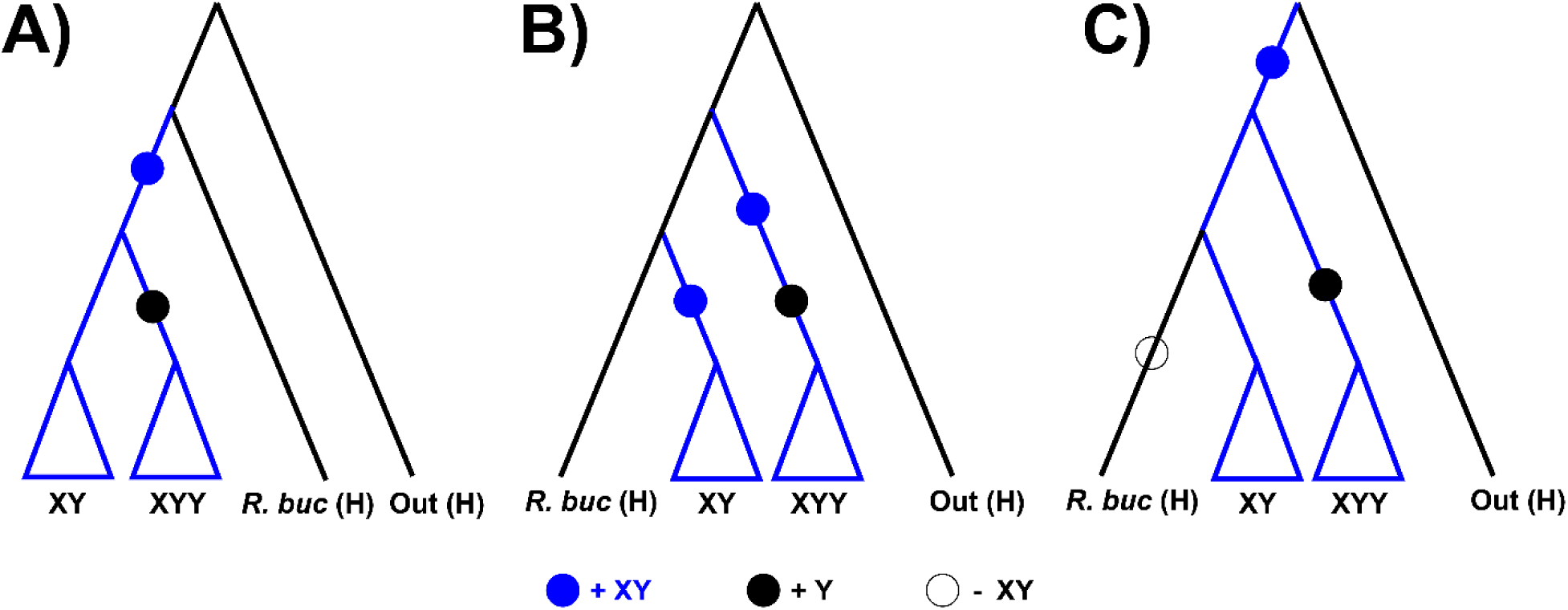
Hypotheses for the evolution and origins of sex chromosomes in *Rumex*. Blue dots indicate the gain of XY sex chromosomes; black dots indicate the gain of an additional Y chromosome; empty dots indicate the loss of XY sex chromosomes. A) the XY and XYY clades are sister, consistent with the phylogeny of Navajas-Pérez et al. (2005) and implying a single origin of the sex chromosomes. B) The XY clade is sister to *R. bucephalophorus* and more distantly related to the XYY clade, consistent with the phylogenies of Grant et al. (2022) and Koenemann et al. (2023). This scenario proposes two independent sex chromosome origins, one in each clade. C) Same phylogeny as scenario B, but now proposing a single sex chromosome origin in the ancestor of the XY clade, XYY clade, and *R. bucephalophorus*, followed by a loss of sex chromosomes in *R. bucephalophorus*.

Here, we present new transcriptome assemblies for 10 *Rumex* species representing the major clades in the genus. We also generate a new high-quality long-read genome assembly for *R. bucephalophorus* and compare genome structure and gene order with assemblies of several additional species in the genus (Sacchi, Humphries et al. 2023). Applying phylogenomic analyses, we find support for two independent origins of sex chromosomes in the genus, consistent with the scenario in Figure 1B. We also find evidence for introgression from unsampled lineages. Lastly, using synteny-based approaches, we find evidence for extensive chromosomal rearrangement in *R. hastatulus* compared to its hermaphroditic relatives. Together, our results highlight the potential of systems like *Rumex* for studying the evolutionary causes and consequences of sex chromosome variation.

## Materials and Methods

### Genomic and transcriptomic data

We conducted RNA-Seq on leaf, bud, and pollen tissue, and assembled the transcriptomes of ten *Rumex* species. In addition to RNA-Seq, we sequenced and assembled the genome of *R. bucephalophorus* using HiFi PacBio sequencing and Dovetail Omni-C sequencing. To complement our new datasets, we obtained recently published genome assemblies of *R. hastatulus*

(Sacchi, Humphries et al. 2023) and Tartary buckwheat, *Fagopyrum tataricum* (Zhang et al. 2017), for a total of 12 species. See the Supplementary Materials and Methods for more detailed sequencing and assembly methods.

### Testing for Whole-Genome Duplications

We used the distribution of *D*_S_ values between paralogs within each of the 12 species to evaluate the presence of whole genome duplications. We first estimated gene trees from each orthogroup codon alignment (see Supplementary section *Orthogroup Identification and Alignment*) using IQ-TREE (Minh et al. 2020a) with the default settings. These gene trees and their corresponding alignments were given to the *codeml* method implemented in PAML (Yang 2007) to estimate values of *D*_N_ and *D*_S_ between each pair of sequences. A custom python script was used to extract *D*_S_ estimates between pairs of sequences belonging to the same species. We then used the R package *mclust* (Scrucca et al. 2016) to evaluate the presence of multiple distributions of log(*D*_S_) values within each species using the Bayesian Information Criterion (BIC). We evaluated the fit of models including 1-9 components, each with equal variance or varying variance. The best-fitting model was selected using the minimum BIC value.

### Phylogenetic Inference

After allowing missing data and single-species duplicates to increase ortholog sampling, we obtained a dataset of 5,263 single-copy genes across 12 species. We used IQ-TREE to estimate a maximum-likelihood phylogeny from a concatenated alignment of all orthologs. To account for the potential effects of incomplete lineage sorting, we also used ASTRAL-III to estimate a summary phylogeny from gene trees estimated for each ortholog. We then time-calibrated our IQ-TREE phylogeny based on fossil evidence and previously estimated node ages (Koenemann et al. 2023). More detailed methods can be found in the Supplementary Materials and Methods.

### Introgression analysis

We tested for introgression among both ancestral and extant lineages using two approaches: the gene tree-based test statistic Δ (Huson et al. 2005, Vanderpool et al. 2020), and a pseudolikelihood approach to estimating phylogenetic networks implemented in the software *PhyloNet* (Than et al. 2008, Yu & Nakhleh 2015). Δ tests for an asymmetry in discordant gene tree counts for a rooted triplet, a classic signature of introgression (Green et al. 2010, Durand et al. 2011). Phylogenetic network estimation is a likelihood-based approach that constructs a network structure containing horizontal branches that denote introgression events. When significant tests involved overlapping sets of taxa, we collapsed them into more ancestral events based on parsimony. More detailed methods can be found in the Supplementary Materials and Methods.

### Resolving the history of sex chromosome evolution

We conducted two analyses to distinguish between two independent origins of sex chromosomes vs. a single origin followed by a loss in *R. bucephalophorus*. First, we BLAST searched a previously generated list of sex-linked genes in *R. rothschildianus* (Crowson et al. 2017) against the genome of XYY *R. hastatulus* to identify shared homologous genes / regions. Second, we estimated gene trees for orthologous genes found in *R. bucephalophorus* and the X and Y chromosomes of *R. hastatulus*, with excess affinity of *R. bucephalophorus* genes to either the X or Y suggesting a potential loss of sex chromosomes. More detailed methods can be found in the Supplementary Materials and Methods.

### Synteny and chromosome-of-origin analyses

Orthology and synteny between protein coding genes in *R. bucephalophorus*, *R. salicifolius*, the XY cytotype of *R. hastatulus* (with a chimeric sex chromosome assembly) and both haplotypes of the *R. hastatulus* XYY cytotype (with phased sex chromosome assemblies) were inferred using GENESPACEv1.1.8 (Lovell et al. 2022). GENESPACE uses MCScanX (Wang et al. 2012) to infer syntenic gene blocks and implements ORTHOFINDERv2.5.4 (Emms and Kelly 2019) and DIAMONDv2.1.4.158 (Buchfink et al. 2021) to find orthogroups within syntenic blocks. Analyses were run and riparian plots visualized in Rv4.1.0. We used the non-default parameter ‘onewayBlast = TRUE’, which is appropriate for species within the same genus, all other parameters were set to default. *Rumex. bucephalophorus* Scaffolds 9 & 10 were excluded from the GENESPACE run as they are very likely to represent separately assembled heterozygous copies of other chromosomes based on the expected chromosome number of 8, and strong similarity of these scaffolds with fragments from other main scaffolds. Scaffolds with fewer than 500 genes were excluded from the plots in all cases.

## Results

### Sequencing and Assembly

We assembled transcriptomes from RNA-Seq data for 10 species that are representative of the major clades of *Rumex* (Supplementary Data 2). Our assemblies were broadly high-quality, with BUSCO-completeness scores ranging from 89% to 95%. The number of main transcripts varied from 23,000 in *R. bucephalophorus* to 56,000 in *R. thyrsiflorus* and was positively correlated with genome size estimates from flow cytometry (Supplementary Data 2). 81% of BUSCO genes were duplicated in *R. thyrsiflorus*; this value ranged from 5% to 16% in other species, potentially suggesting a large-scale gene duplication or whole-genome duplication event (see Discussion).

We additionally generated a high-quality chromosome-scale assembly of *R. bucephalophorus* using high-coverage HiFi PAC Bio sequencing and Dovetail Omni-C sequencing (Supplementary Figure 1, Supplementary Table 2). After removing erroneously separately assembled chromosome haplotypes (see Supplementary Materials and Methods), the assembly size is 2.062 GB (compared to flow cytometry estimates of 1.96 GB), with 88% of the genome found in the main 8 scaffolds, consistent with karyotypic evidence for eight autosomes in the species (Navajas-Pérez et al. 2005).

### Mixed evidence for recent whole-genome duplications

Previous work has identified an ancient whole-genome duplication (WGD) shared by buckwheat and *Rumex* (Zhang et al. 2017, Fawcett et al. 2023). In addition, *R. acetosella*, *R. scutatus* and *R. paucifolius* are known to have natural polyploid populations (Löve 1940, Löve 1942, Smith 1968). To further assess the presence of recent polyploidy events in our dataset, we calculated dS values between gene paralogs for each species. In the absence of whole-genome duplications, dS values should be exponentially distributed (normally distributed in log-space) following a birth-death model for gene gain and loss (Lynch & Conery 2000, Blanc & Wolfe 2004). WGDs introduce numerous gene duplications at the same point in time, which should result in additional peaks in the distribution of dS values between paralogs.

We found mixed evidence for recent whole-genome duplications among our study species. (Figure 2, Supplementary Figure 4). There is equivocal statistical support (based on minimum BIC value) for one or two distributions of dS values in *R. acetosella*, *R. paucifolius*, and *R. thyrsiflorus*, and support for multiple distributions in *R. bucephalophorus* (Supplementary Figure 4). *R. acetosella* and *R. paucifolius*, two known polyploid species, share a peak of log(*dS*) values at approximately –2.5 (Figure 2), corresponding to a *dS* value of ∼0.08. BIC values for *R. bucephalophorus* support up to eight distributions of *dS* values (Supplementary Figure 4); given the low number of paralogs sampled for this species, and the lack of observed polyploids in previous studies, this is likely technical error. Finally, we observe a large peak of *dS* values in *R. thyrsiflorus* at approximately –1, corresponding to a *dS* value of ∼0.35. No known polyploids have been observed in this species. Overall, while we identify mixed evidence for recent whole genome duplication events, our approach to identification of single-copy orthologues while allowing for species-specific duplications should enable a robust phylogeny even in the presence of some autopolyploid lineages (however, an allopolyploid lineage could lead to inference challenges; see Discussion).

**Figure 2:**
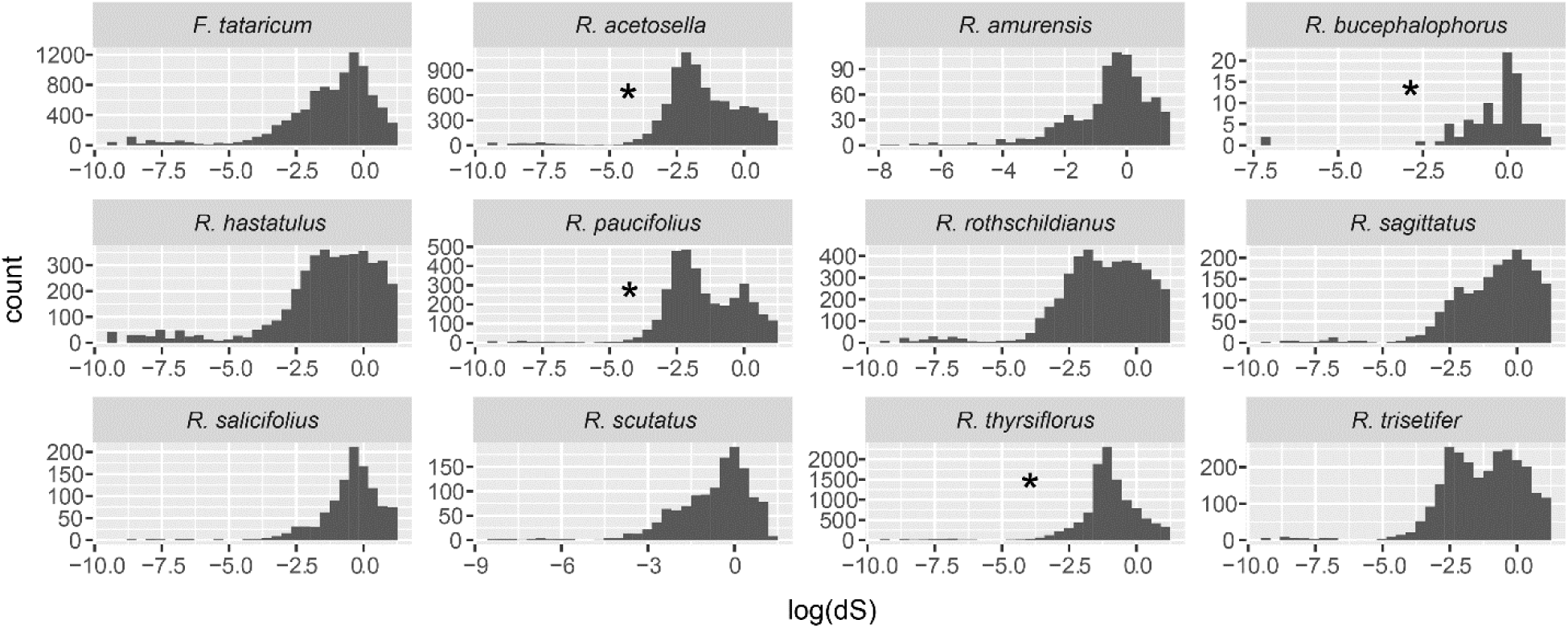
Distribution of log(*dS*) values between gene paralogs for each of our 12 studied *Rumex* species. Species with support for multiple distributions of values are marked with an asterisk.

### Whole-transcriptome phylogeny supports R. bucephalophorus as sister to the XY clade

We used our 12-species transcriptomic/genomic dataset to infer a phylogeny of *Rumex* using both concatenated maximum-likelihood (IQ-TREE) and gene-tree summary (ASTRAL-III) approaches. Our maximum-likelihood tree was estimated with strong statistical support, with all nodes having 100% support in both SH-aLRT and ultrafast bootstrap measures (Figure 3). The topology is in general agreement with previous studies, supporting two monophyletic clades with sex chromosomes and two earlier-diverging hermaphroditic clades. We place *R. bucephalophorus* as sister to the XY clade, to the exclusion of the more distant XYY clade, in agreement with the chloroplast phylogenies of Grant et al. (2022) and Koenemann et al. (2023), but in contrast to the phylogeny of Navajas-Pérez et al. (2005). We also infer a sister relationship between *R. hastatulus* and *R. paucifolius*; this inference agrees with Koenemann et al. (2023), but contrasts with Navajas-Pérez et al. (2005) and Grant et al. (2022), who place *R. hastatulus* as sister to *R. acetosella*. Our divergence time estimates generally agree with those of Koenemann et al. (2023), though we infer an older node age for the XY clade (5.6 MYA vs. 2.61 MYA), and a more recent node age for the root (16 MYA vs. 22.13 MYA).

**Figure 3:**
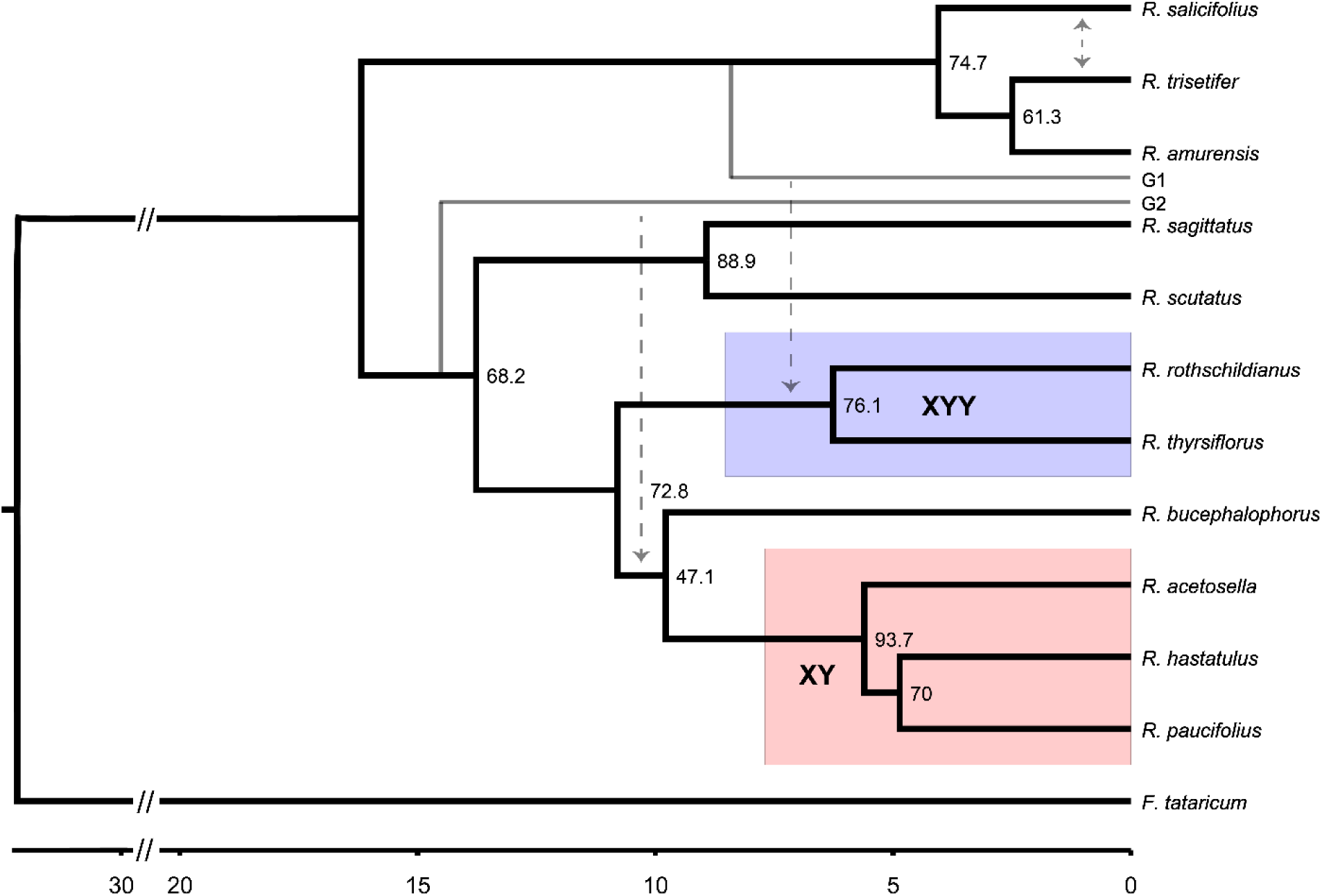
Whole-transcriptome maximum-likelihood phylogeny of 11 *Rumex* species, with branch length units in millions of years (x-axis scale). Branch length distance to the root is truncated for visual clarity. The XY and XYY clades are highlighted in red and blue, respectively. Nodes are labelled with gene concordance factors. Grey dashed arrows indicate inferred introgression events. Unsampled lineages involved in introgression events are denoted with the grey branches labelled “G1” and “G2”.

Gene tree discordance varied among clades but was not prevalent enough to generate substantial phylogenetic uncertainty. Highlighting this, our ASTRAL-III phylogeny returned the same topology as maximum-likelihood (Supplementary Figure 5). As a gene-tree summary approach, ASTRAL-III is more robust to high rates of incomplete lineage sorting that can mislead standard ML approaches (Mirarab et al. 2014). Gene concordance factors (gCFs), a measure of the proportion of gene trees in the dataset consistent with each branch, varied from 47.1% to 93.7% (Figure 3, Supplementary Table 3). The lowest gCF was at the node where *R. bucephalophorus* splits from the ancestor of the XY clade, at 47.1%. This finding helps explain the uncertainty in its placement in previous studies. In contrast, most branches in the phylogeny exhibit modest levels of discordance, being supported by between 60% and 94% of gene trees (Figure 3).

### Signatures of ghost introgression in the Rumex phylogeny

We investigated signatures of introgression among our sampled species using a test statistic, Δ, based on gene tree counts, in addition to inferring phylogenetic networks with *PhyloNet*. Our Δ tests returned a multitude of highly significant results, often implying introgression between lineages that were not contemporaneous (according to the phylogeny of Figure 3) (Supplementary Data 3). On further examination of our results, we observed that many species, when included in one of the sister species positions in a test, often implied the other two species in the test as introgressing with each other, regardless of their identity. For instance, in the triplet [(A,X),Y], where A is a particular species and X and Y could be any two species with the specified relationship, introgression would always be implied between X and Y. This indicates that X is more distantly related to A than expected based on phylogenetic relationships, a classic signature of ghost introgression from an earlier-diverging donor lineage (Supplementary Figure 6) (Ottenburghs 2020, Tricou et al. 2022a, Tricou et al. 2022b). Such ghost introgression events might be expected in our study because we have sampled only 11 of the 200 described species in the genus. These unexpectedly distant species include *R. thyrsiflorus*, *R. rothschildianus, R. acetosella*, *R. hastatulus*, *R. paucifolius*, and *R. bucephalophorus*.

Our best-fitting phylogenetic network supports the existence of two ghost introgression events but disagrees with our inferred species tree topology in several places (Supplementary Figure 7). As our phylogeny has strong statistical and genealogical support, we chose to reconcile the phylogeny with our best-fitting phylogenetic network and set of significant Δ statistics to propose two ghost introgression events (Figure 3). The first event was into the common ancestor of the clade containing *R. bucephalophorus* and the XY species. The donor lineage for this event is likely an early-diverging member of the clade containing the two sex chromosome subclades (branch “G2” in Figure 3), possibly *Rumex induratus* or a close relative based on the phylogeny of Grant et al. (2022). The other event involved the ancestor of *R. thyrsiflorus* and *R. rothschildianus*, with the donor likely a member of the early-diverging hermaphroditic clade containing *R. salicifolius* and its relatives (branch “G1” in Figure 3). Finally, we see additional evidence of introgression between *R. salicifolius* and *R. trisetifer* (Figure 3), two closely related hermaphroditic species.

### Independent evolution of XY sex chromosomes

Our updated phylogeny of *Rumex* rules out a simple single-origin scenario (Figure 1A) for the evolution of sex chromosomes. We conducted additional analyses to distinguish between the two remaining possibilities: two independent origins of sex chromosomes (Figure 1B) vs. a single origin followed by loss of sex chromosomes in *R. bucephalophorus* (Figure 1C). First, we evaluated the chromosome of origin of sex-linked genes in the two major sex chromosome clades by mapping previous transcriptome-identified sex-linked genes from *R. rothchildianus* (XYY clade) to our genome assembly of *R. hastatulus* (XY clade). In the simplest scenario, where a fully-formed sex chromosome evolves once and is inherited by both groups, we would expect sex-linked genes in *R. rothschildianus* to map primarily to the X chromosome of *R. hastatulus*. Alternatively, if sex chromosomes originated or evolved independently in the two groups, sex-linked genes in *R. rothschildianus* should map to some combination of autosomes and/or the X chromosome. We found that sex-linked genes in *R. rothschildianus* mapped to all chromosomes of *R. hastatulus*, with most mapping to autosomes 1 and 2, followed by the X chromosome, and a small number of hits on autosome 4 (Figure 4, left column).

**Figure 4:**
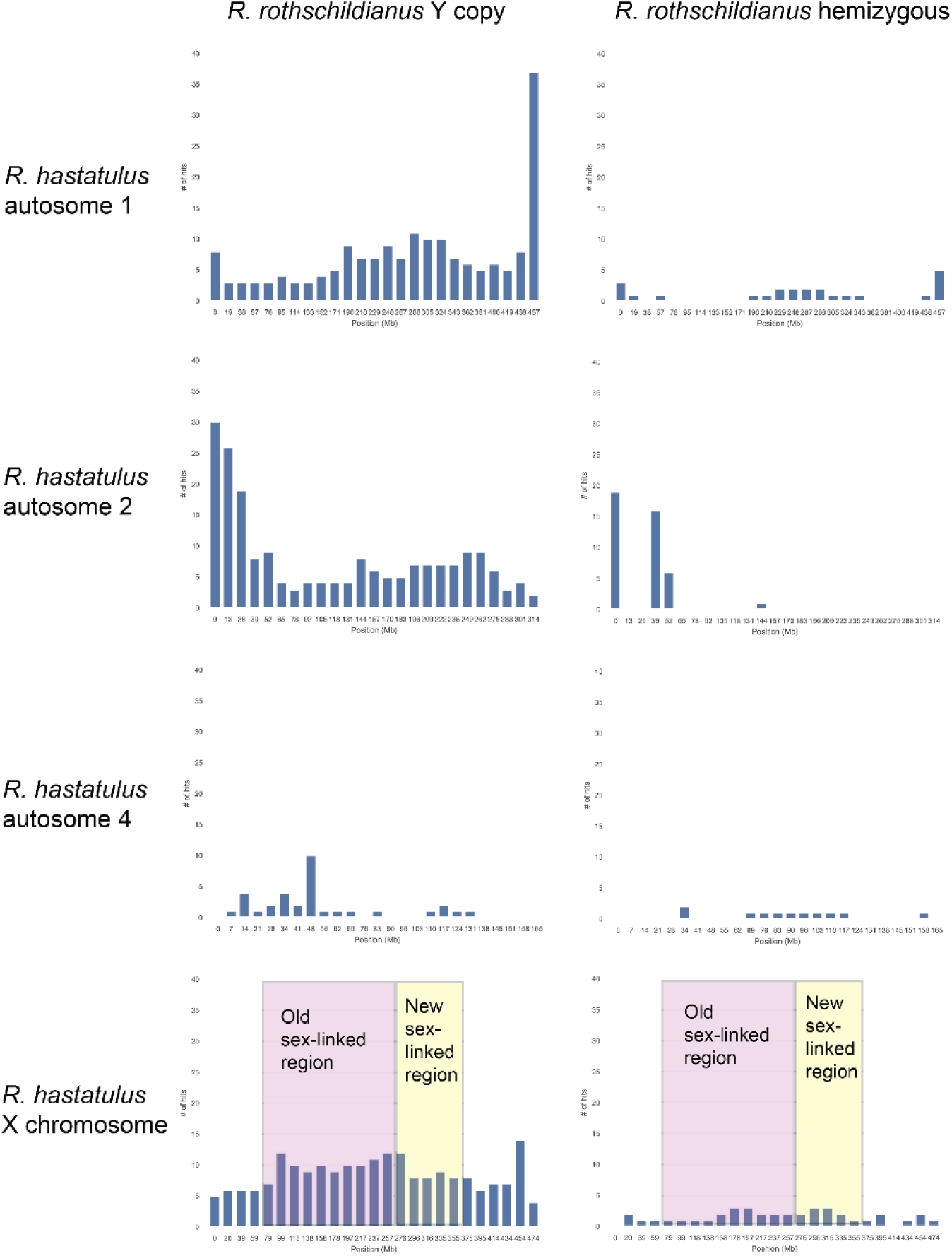
Distribution of BLAST hits of sex-linked genes in *Rumex rothschildianus* against the genome of *R. hastatulus*, divided into 25 equally spaced windows. In each plot, the x-axis is the position on the chromosome in megabases (Mb) at the start of the window, and the y-axis is the number of BLAST hits found within that window. Left column is X-linked genes in *R. rothschildianus* where a Y copy is still present; right column is hemizygous X-linked genes. Each row shows results for a chromosome in *R. hastatulus*. In the *R. hastatulus* X chromosome, the approximate location of the old sex-linked region shared by both cytotypes is highlighted in magenta, and the new sex-linked region in the XYY cytotype is highlighted in yellow. Remaining regions of the X chromosome represent pseudoautosomal regions in *R. hastatulus*.

We observed a significant number of hits on the *R. hastatulus* X chromosome, raising the possibility that the two sex chromosome clades share a “core” sex-determining region with a single origin, with chromosomal rearrangements within the two clades resulting in independent origins for other regions of the X. To evaluate this hypothesis further, we separately examined chromosome-of-origin for hemizygous genes and genes with a Y copy present in *R. rothschildianus*. Hemizygous genes are generally expected to be older and would therefore more likely be found in an older shared sex-determining region, whereas genes with a Y copy are expected to be younger and therefore more common in younger, independently evolving regions of the sex chromosomes. In this case, most genes mapped to a region containing the first 50 MB of autosome 2, with smaller numbers of genes distributed across the remaining chromosomes (Figure 4, right column).

We found a slight elevation in the number of genes mapped to the sex-linked regions of the *R. hastatulus* X (Figure 4, bottom row). Otherwise, the distribution of genes mapped to *R. hastatulus* broadly corresponds to the overall density and number of genes along each chromosome (Rifkin et al. 2022, Sacchi, Humphries et al. 2023). One explanation for this pattern is that extensive chromosomal rearrangements in *R. hastatulus* (see results section *Elevated rates of chromosomal rearrangement in R. hastatulus*, Figure 6) have largely randomized the locations of genes with respect to their ancestral homologs, resulting in BLAST hits resembling a random draw of genes from the chromosome. Overall, these results are most consistent with the independent evolution of the sex chromosomes, though we cannot rule out the single origin of a smaller sex-determining region shared by the two groups.

### No ancestral sex chromosome system in R. bucephalophorus

To further resolve the history of sex chromosome evolution, we next examined the relationship of *R. bucephalophorus* genes to orthologous X/Y gametologs present in *R. hastatulus*. Evolutionary loss of sex chromosomes is generally expected to proceed via an inactivating mutation on one of the sex chromosomes that restores sex-specific functions (e.g. Vicoso & Bachtrog 2013) and in this case allows for the production of hermaphroditic individuals. The mutated sex chromosome then becomes an autosome, while the other is lost from the population. Therefore, if *R. bucephalophorus* lost an XY system that is shared by the two extant XY clades, formerly sex-linked genes should coalesce primarily with either the X chromosome (in the case of loss of the Y) or Y chromosome (in case of loss of the X) of *R. hastatulus* (Figure 5A). Alternatively, if the sex chromosomes arose independently in the XY and XYY clades, and *R. bucephalophorus* has simply retained the ancestral hermaphroditic state, then XY gametologs in *R. hastatulus* should coalesce with each other before their ortholog in *R. bucephalophorus* (Figure 5A), consistent with the phylogeny. This sets up a symmetric expectation, where a loss of sex chromosomes in *R. bucephalophorus* should lead to most trees having one of the two possible discordant histories (Figure 5A).

**Figure 5:**
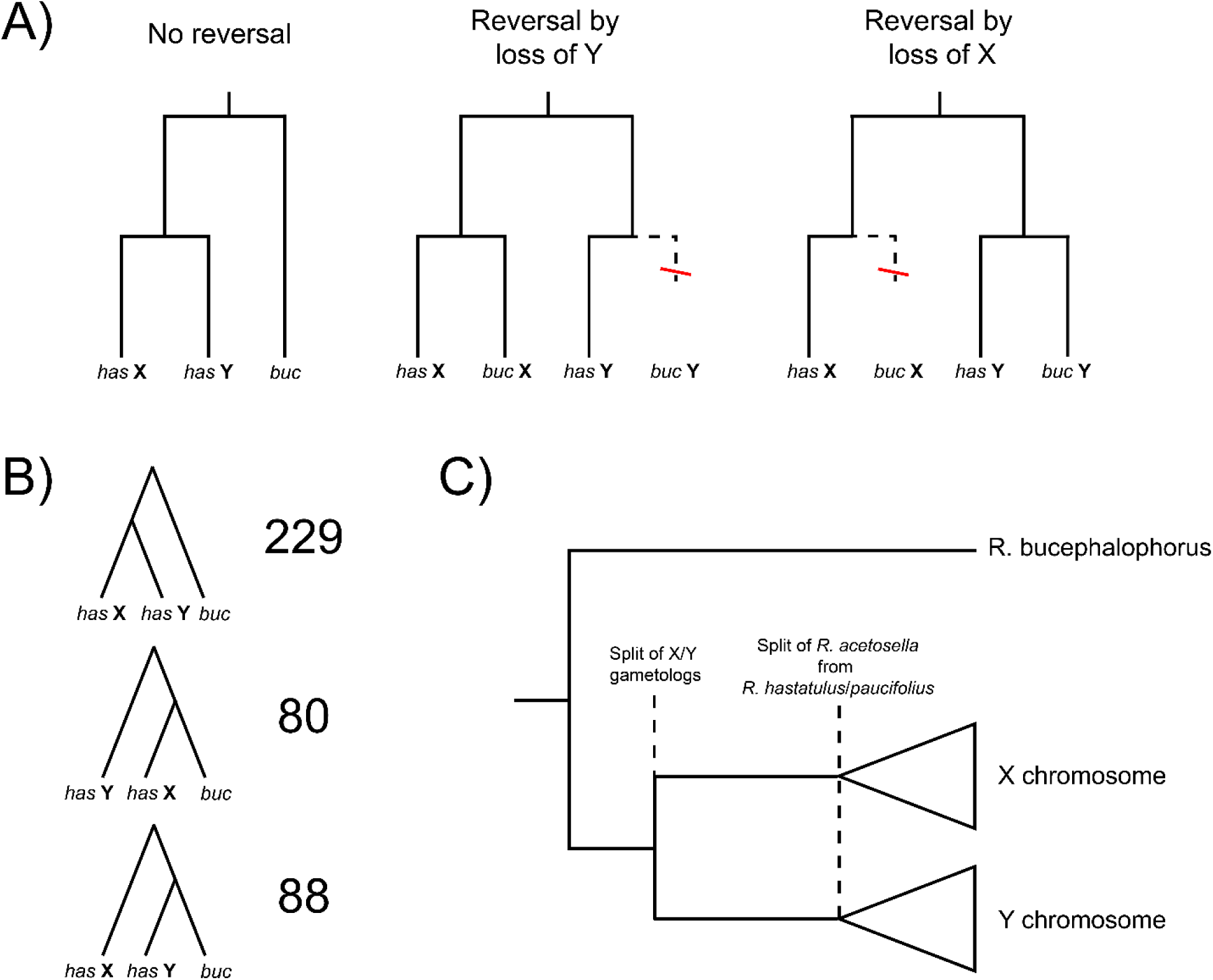
Relationship of *Rumex hastatulus* X/Y gametologs to *R. bucephalophorus*. A) Three scenarios for the evolution of sex chromosomes in *R. bucephalophorus*. For each scenario, the expected majority gene tree is traced by the solid black line. B) Counts of the three possible tree topologies in coding sequences shared by *R. bucephalophorus* and the X and Y chromosomes of *R. hastatulus*. C) Demographic history explaining our observed gene tree counts in panel B. Recombination suppression between the X and Y arises relatively quickly in the ancestor of the X/Y clade after its split from *R. bucephalophorus*. Subsequent speciation within the XY clade happens later on.

Out of 397 single-copy genes present in the old sex-linked region of the *R. hastatulus* X and Y chromosomes and *R. bucephalophorus*, 229 (57.7%) support the species phylogeny, 80 (20.1%) support the loss of Y scenario, and 88 (22.2%) support the loss of X scenario (Figure 5B). This suggests a historical absence of sex chromosomes in *R. bucephalophorus*; therefore, our results support two independent gains of sex chromosomes. Interestingly, the proportion of discordant topologies (42.3%) is much larger than the genome-wide average, where 94% of gene trees support monophyly of the XY clade (Figure 3). One possible explanation for this pattern is that the sex chromosomes evolved relatively quickly in the common ancestor of the XY clade after its split from *R. bucephalophorus* (Figure 5C). The short amount of time separating this split from the divergence of X/Y gametologs would result in higher rates of incomplete lineage sorting (Figure 5C), producing the two discordant histories with equal frequency. Consistent with this explanation, Crowson et al. (2017) estimated the timing of initial recombination suppression in the *R. hastatulus* X chromosome at 9-16 million years ago, much older than our age estimate for the XY clade of 5.6 MYA (Figure 3). The more recent end of this estimate at 9MYA falls after the split of *R. bucephalophorus* at approximately 10 MYA (Figure 3), suggesting these events could plausibly have occurred in relatively short succession.

**Figure 6:**
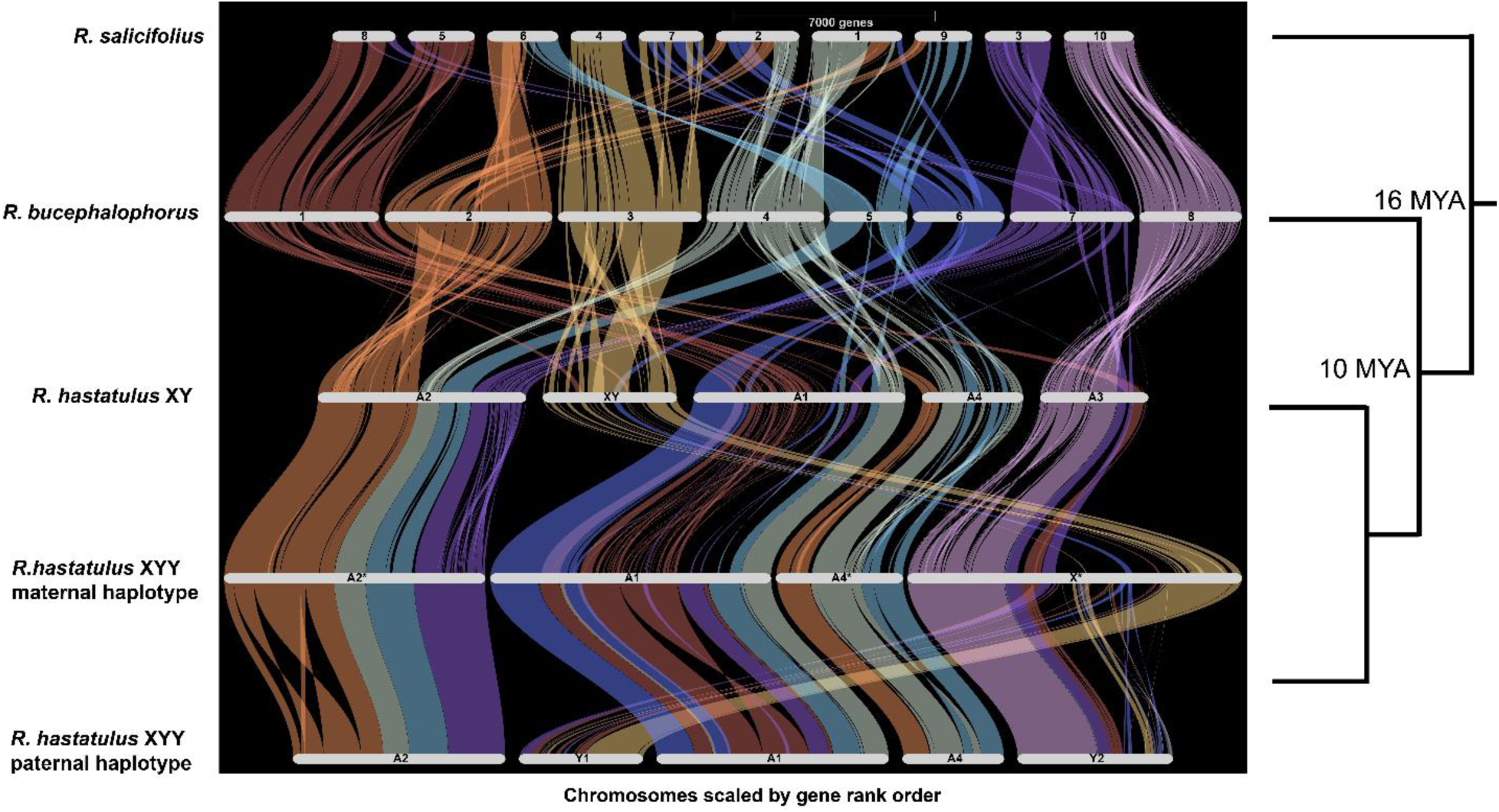
GENESPACE riparian plots showing synteny between *Rumex salificolius*, *R. bucephalophorus*, and the two cytotypes of *R. hastatulus*. Syntenic blocks are ordered and colored according to chromosome of origin in *R. bucephalophorus*. Branch lengths on righthand tree are not to scale. Note that chromosome size is scaled by gene rank order, rather than by physical size.

### Elevated rates of chromosomal rearrangement in R. hastatulus

To futher investigate the origin and evolution of sex chromosomes in *R. hastatulus*, we sought to identify the ancestral autosomal homologs of the sex chromosomes in the XY clade and investigate the history of rearrangements among these chromosomes. We conducted synteny analysis on genome assemblies of *R. salicifolius* (Sacchi, Humphries et al. 2023), both cytotypes of *R. hastatulus* (Sacchi, Humphries et al. 2023), and our de novo assembly of *R. bucephalophorus* using the software GENESPACE (Lovell et al. 2022) (Figure 6), suggesting a relatively simple history of chromosomal fusion. Although there are some karyotypic differences (*R. salificolius* has 10 chromosome pairs and *R. bucephalophorus* has 8), the genomes of the two hermaphroditic species are largely collinear; most chromosomes in *R. salificolius* are syntentic with one or two chromosomal segments (likely representing chromosome arms) in *R. bucephalophorus* (Figure 6). In contrast, despite being more closely related to *R. bucephalophorus*, the genome of *R. hastatulus* is highly rearranged. In addition to a further reduction in chromosome number (with four autosomes plus the sex chromosome pair), each chromosome of *R. hastatulus* is syntenic with 3-6 chromosomal segments of *R. bucephalophorus* (Figure 6). These results suggest an elevated rate of chromosomal rearrangement across the genome of *R. hastatulus*, or possibly the XY clade at large (though additional genome sequencing of *R. acetosella/paucifolius* would be needed to confirm this possibility). *Rumex acetosella* and *R. paucifolius* both have reduced chromosome number (*n*=7) relative to *R. bucephalophorus* (*n*=8) (Navajas-Pérez et al. 2005), so it is likely that at least some of this rearrangement is ancestral to the XY clade.

The sex chromosomes of *R. hastatulus* have complex chromosomal origins, with key regions sharing syntenic blocks with different chromosomes in hermaphroditic relatives (Figure 7). The old sex-linked region shared by both cytotypes (dark blue in Fig. 7) is orthologous primarily with chromosome 3 of *R. bucephalophorus*, with smaller syntenic blocks on chromosomes 1 and 6; the PAR for this region (light blue, Fig. 7) is also orthologous to chromosome 3. The neo-sex-linked region (yellow, Fig. 7) contains syntenic blocks from chromosomes 1 and 7 of *R. bucephalophorus*, while the neo-PAR (light red, Fig. 7) is orthologous with chromosome 8. These three syntenic blocks are all present on autosome 3 of XY *R. hastatulus*, suggesting their fusion predates the sex chromosome-autosome fusion that formed the XYY cytotype. A relatively small central region of chromosome 1 in *R. bucephalophorus* has independently contributed syntenic blocks to both the old and neo-X chromosomes of *R. hastatulus*, identifying this as an interesting region for future study. Finally, a reciprocal translocation of the two Y chromosomes in the XYY cytotype has resulted in both old and neo-X syntenic blocks existing in both chromosomes (Figure 7, Sacchi, Humphries et al. 2023).

**Figure 7:**
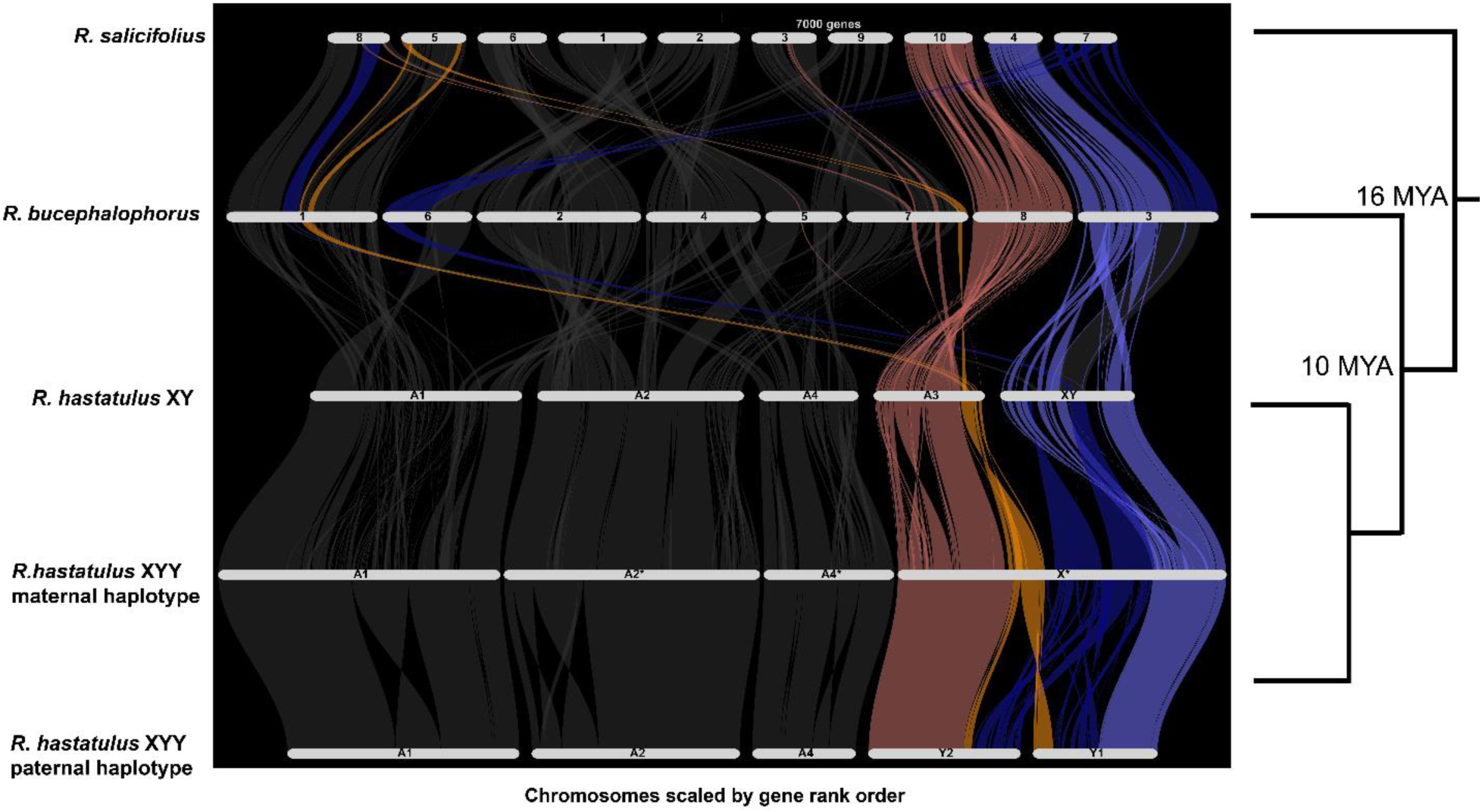
GENESPACE riparian plots showing synteny between *Rumex salificolius*, *R. bucephalophorus*, and the sex chromosomes of *R. hastatulus*. Syntenic blocks are ordered and colored according to regions of the neo-X chromosome of *R. hastatulus*: neo-PAR in light red, neo-sex-linked region in yellow, old sex-linked region in dark blue, and old PAR in light blue. Branch lengths on righthand tree are not to scale. Note that chromosome size is scaled by gene rank order, rather than physical size.

## Discussion

Phylogenetic inference must often compromise between taxon sampling and sampling of genetic loci due to computational and sampling constraints. Previous studies estimating phylogenies of *Rumex* focused on sampling a wide range of taxa, at the cost of using a relatively small number (1-3) of genetic markers (Navajas-Pérez et al. 2005, Grant et al. 2022, Koenemann et al. 2023). Whereas increased taxon sampling can improve statistical confidence in inferred relationships, it will not help if particular nodes are incorrectly resolved due to biological sources of gene tree discordance such as incomplete lineage sorting (Degnan & Rosenberg 2009). Here, we used genome-scale sampling of loci with a smaller set of taxa to resolve key relationships within *Rumex*, with important implications for the history of sex chromosome evolution in the genus. We found that the node where *R. bucephalophorus* branches off is supported by fewer than half of our estimated gene trees (Figure 3), highlighting the need to sample many loci to accurately resolve relationships.

Our greater sampling of loci also allowed us to test for introgression, which cannot be done with a small set of genetic markers. Introgression involving unsampled taxa (“ghost” introgression) is increasingly recognized as an issue for phylogenetic inference (Ottenburghs 2020, Tricou et al. 2022a, Tricou et al. 2022b), but it is still challenging to estimate from genomic data as it is easily confused with introgression among sampled taxa. Inference requires careful examination of the three-species introgression results and patterns in genomic data, as we have done here. As highlighted by our results and recent simulation work (Tricou et al. 2022a, Tricou et al. 2022b), authors should explicitly consider ghost introgression as a parsimonious hypotheses when studying introgression. We should note, however, that such parsimony-based inferences might eliminate true instances of introgression among sampled taxa, including our study; full-likelihood methods with more explicit model selection criteria like the PhyloNet method we applied in our study can further aid in distinguishing among scenarios. Although we do not know the identity of the donor lineages, the timing of our proposed events likely predates the evolution of sex chromosomes in both clades (Figure 3). Additionally, the one instance of more recent introgression we observe is between two closely related hermaphroditic species. While preliminary, our results are consistent with the idea that sex chromosome and sexual system differences among *Rumex* species form significant barriers to more recent introgression.

We applied mixture model analyses to the distribution of *dS* values in each of our study species to identify whole-genome duplications. This kind of analysis has important limitations; it has a tendency to over-estimate the number of WGD events and has poor power when the number of retained gene duplicates is low (Tiley et al. 2018). These limitations are compounded when using transcriptomic data, as pseudogenized duplicate gene copies that are no longer expressed cannot be detected. Nonetheless, we found mixed evidence for two distributions of *dS* values in two species known to have both polyploid and diploid varieties, *R. acetosella* and *R. paucifolius* (Supplementary Data 2, Figure 2, Supplementary Figure 4). These species share a peak of *dS* values at approximately 0.08, corresponding to approximately 5 million years of divergence (following calculations in Crowson et al. 2017). The age and shared peak imply the WGD may have occurred in an ancestral population of the XY clade, and the polyploid cytotype could have subsequently been lost in *R. hastatulus*. The polyploid variety could also have re-evolved diploidy, which would explain the extensive genomic rearrangement we observed in *R. hastatulus*. However, it is unlikely that the signal of such an event would be completely erased after only 5 million years (Li et al. 2021).

The large peak we observed in *R. thyrsiflorus* is more difficult to explain, as it is much older (*dS* ∼0.35) and not shared by other species, and polyploidy has not been observed in the species. While old, the peak is too young to be explained by differential retention of the ancient WGD shared with buckwheat, which occurred ∼70 MYA (Fawcett et al. 2023). One potential explanation is a large burst of gene duplication without polyploidy; a large genome size and high percentage of duplicated BUSCO genes in our transcriptome assembly support this idea (Supplementary Data 2). It is also possible that we have identified previously unknown polyploidy in the species; however, polyploidy should not be tolerated under the X-autosome balance mechanism of sex determination expected to be used by members of the XYY clade such as *R. thyrsiflorus* and its relatives (Mable 2004). In general, because we sample single gene copies and limit gene duplications to a single species, our phylogenomic analyses should be robust to the effects of autopolyploidy across the genus, regardless of the precise history of events. However, allopolyploidy would complicate our analysis, as there would be substantial disagreement in genealogical relationships in the genome of the allopolyploid lineage. Ultimately, whole-genome sequences will be required to fully resolve the history of WGD in the genus.

We found that previously identified sex-linked genes in *R. rothschildianus* are homologous to all autosomes and the X chromosome (both old X and neo-X) of *R. hastatulus*, with a small number of hits on autosome 4 (Figure 4). While this result adds some evidence of shared sex-linked genes between these species (in contrast with the previous transcriptome-based results of Crowson et al. 2017), the large number of sex-linked genes from *R. hastatulus* autosomes provide clear support for significantly independent evolution of sex chromosomes in the two major clades. Because of the additional independent evolution of an XYY system (likely from an XY ancestor) in the clade with *R. rothschidianus*, we expected a priori some differences in patterns of sex linkage. The number of genes orthologous to each chromosome, as well as their concentration at the ends of the chromosomes, is broadly consistent with overall patterns of gene density in *R. hastatulus* (Rifkin et al. 2022, Sacchi, Humphries et al. 2023). This suggests a largely random distribution of genes, which could arise from the high rate of chromosome rearrangement we observed (Figure 6). Nonetheless, there are some interesting deviations; many genes mapped to the last 10 MB of autosome 1 and the first 50 MB of autosome 2. Both regions could simply be preserved syntenic blocks with ancestrally high gene content, but it is interesting that hemizygous *R. rothschildianus* genes appear to be particularly enriched at the beginning of autosome 2. We also found a slight elevation of genes mapping to the sex-linked regions of the X chromosome (Figure 4), despite this region having relatively low gene density overall. This result could be explained by recruitment of genes with ancestrally sex-biased functions to the X chromosome, or potentially by a shared sex-determining locus between the two major clades. Ultimately, chromosome-scale genome assemblies of *R. rothschildianus* and close relatives of *R. hastatulus* will be needed to further resolve the complex history of karyotypic evolution in the genus.

Although our combined results support the largely independent origins of the sex chromosomes in the two major dioecious clades, this does not fully rule out a single origin of dioecy. It is possible that the two major sex chromosome clades originally shared a sex-determining region that arose in their common ancestor. Alternatively, introgression between the XYY clade and the XY clade following its split with *R. bucephalophorus* could have led to a shared genetic basis of SD. In either case, subsequent rearrangements and a divergent history of recombination suppression would drive highly divergent sex-linkage of many genes in the two groups, including an ancient X-autosome fusion that gave rise to the XYY karyotype. While we find no evidence for loss of sex chromosomes in *R. bucephalophorus*, incomplete lineage sorting of the sex-determining locus could have led to its inheritance in the two major sex chromosome clades, but not *R. bucephalophorus*, following a single origin in their common ancestor (Avise & Robinson 2008, Mendes & Hahn 2016). Our introgression analyses do not provide clear support for introgression among these clades, but as previously mentioned, our results made it challenging to distinguish between ancient introgression among unsampled hermaphroditic lineages and more contemporary bouts of introgression among sampled taxa. Identification of the causal sex-determining genes across the genus would enable a direct examination of these possibilities further. However, the evidence for divergent mechanisms across species means that the two clades are unlikely to have a single genetic basis currently, even if there was original sharing of the mechanism of sex determination.

Classic theory predicts an elevated rate of fusions involving sex chromosomes and autosomes, which helps physically link sexually antagonistic variation on other chromosomes to the sex-determining region (Charlesworth & Charlesworth 1980, Rice 1987, Charlesworth et al. 2005). Chromosomal rearrangements including fusions are frequent in *Rumex*, as evidenced by successive reductions in chromosome number from the ancestral x=10 karyotype (Navajas-Pérez et al. 2005), high rates of intraspecific rearrangement in XYY species *R. acetosa* (Parker & Wilby 1989), and our synteny analyses (Figures 5 and 6). Although we can confirm the existence of an X-autosome fusion forming a neo-X chromosome in *R. hastatulus* (Figure 7), the rate of rearrangements involving sex chromosomes does not appear to be elevated relative to autosomes in this species. Autosome 1 and the neo-X chromosome are both syntenic to six chromosomal regions of *R. bucephalophorus*, for example. Given that rates of rearrangement are generally elevated in *R. hastatulus*, we do not need to invoke an adaptive explanation related to sexually antagonistic selection for the number of fusions observed on the sex chromosomes. On the other hand, it may be that the “baseline” rate of rearrangement (either due to adaptation (Guerrero & Kirkpatrick 2014) or a higher rate of mutation) is sufficient to capture SA variation on the sex chromosomes, without driving an elevated rate. Depending on the sequence of events, this elevated rate of rearrangement may have even promoted the formation of sex chromosomes in the first place by allowing SA variation to be captured shortly after the origins of a sex-determining locus. Genome sequencing of close relatives of *R. hastatulus* will be necessary to determine if this elevated rate of rearrangement is a feature specific to *R. hastatulus* or is common to dioecious species in general.

Our study focused primarily on the sex chromosomes, but even among species without them, *Rumex* contains a variety of sexual systems. Among our studied species without differentiated sex chromosomes, *R. bucephalophorus* has been observed in our samples and described in other studies as gynomonoecious (female and bisexual flowers in the same individual) (Talavera et al. 2011), *R. sagittatus* has been described as dioecious / monoecious but without heteromorphic sex chromosomes (male and female flowers in the same individual, Navajas-Pérez et al. 2005) (though our samples were hermaphroditic), and *R. scutatus* has been described as polygamous (male, female, and bisexual flowers in the same individual, Navajas-Pérez et al. 2005). Hermaphroditic individuals have also been described for *R. bucephalophorus* and *R. scutatus*. This variation is important because the evolutionary transition from hermaphroditism to dioecy is expected to proceed through these “intermediate” sexual systems (Barrett 2002). The most important of these pathways is thought to be through gynodioecy (female and bisexual flowers in different individuals), via the successive fixation of male-inactivating and female-inactivating mutations producing separate male and female individuals (Charlesworth & Charlesworth 1978, Spigler & Ashman 2012). Gynodioecy is not observed among our study species, although it has been described in members of the clade that includes *R. sagittatus* and *R. scutatus* (Navajas-Pérez et al. 2005). Alternatively, dioecy could arise from gynomonoecy via disruptive selection for increased investment into male reproduction in bisexual flowers (Barrett 2002). Unfortunately, given the wide variation of sexual systems in our study species and our lower taxon sampling, we have insufficient information in our study to reconstruct the ancestral sexual system to the two sex chromosome clades. Regardless, it is clear that significant ancestral variation in sexual systems would have existed to facilitate transitions from hermaphroditism to dioecy across the genus.

Our work opens many directions for future research. First, classic models of sex chromosome evolution predict the progressive suppression of recombination along the length of the chromosome, forming evolutionary strata. Our genome assembly of *R. bucephalophorus*, in addition to recently generated assemblies of *R. hastatulus* (Sacchi, Humphries et al. 2023) and hermaphroditic relatives, will allow for investigations into the evolution of recombination suppression along the sex chromosomes of *R. hastatulus*. Second, our finding of independent origins of sex chromosomes raises interesting questions about the XYY clade. Overlap in the ancestral autosomal homologs contributing to these two sex chromosome systems, including the formation of the neo-X and extra Y chromosome in the XYY clade, could hint at ancestral sexually antagonistic variation promoting the formation of sex chromosomes in the genus. Answering these questions will require genome assemblies from representatives of the XYY clade. Finally, our transcriptome assemblies and estimated phylogeny have great potential to address questions related to the evolution of gene expression. Phylogenetic frameworks will allow future studies to understand the role that sex-biased gene expression and dosage compensation play in driving or being driven by the evolution of sex chromosomes. Increased taxon sampling from this species-rich genus within a phylogenomic framework will increase the power to address all these questions. Overall, our study opens the door to *Rumex* as an exciting system for the comparative study of sex chromosome evolution.

### Data Availability

Genome and transcriptome assemblies will be available at COGE (https://genomevolution.org/coge/) at xxxx and GenBank at xxxx. Raw sequencing reads for RNA-Seq are available at the SRA under BioProject PRJNA698922, and for genome sequencing under XYZ. Customs scripts and supplementary data files are available on GitHub at https://github.com/mhibbins/RumexComparative.

## Acknowledgements

We thank Bill Cole, Thomas Gludovacz, Emily Glasgow, Madeline Jarvis-Cross, and Deanna Kim for help with plant growth and maintenance, and Matthew Hahn for comments on the manuscript. This research was supported by NSERC Discovery Grants awarded to SCHB and SIW, and an EEB Postdoctoral Fellowship awarded to MSH by the EEB Department at the University of Toronto.

## Supplementary Materials and Methods

### Sample Collection and Sequencing

For RNASeq, samples for live tissue were sourced from the USDA GRIN network (USDA Agricultural Research Service), the Southwest China Wildlife Germplasm Genobank (http://www.genobank.org/), and the collections from Spencer Barrett’s lab, totaling 10 species (see Supplementary Table 1). Plants were grown under glasshouse conditions between 2018 and 2020. Because many *Rumex* species are perennials, we collected leaf, bud, and pollen tissue opportunistically based on availability. Leaf and bud tissue were collected directly into LN2 using sterilized forceps. Pollen was collected using keif boxes (WACKY WILLYS,Inc. BC, Canada) and either frozen in LN2 or germinated prior to sequencing using the medium developed in Adhikari & Campbell (1998).

For the genome assembly of *R. bucephalophorus*, we collected open-pollinated seeds from a population at Poleg, Netanya, Israel in March 2019. We placed the seeds on moist filter paper in a petri dish at 4 °C for at least 24 hours, then left the petri dish at room temperature. After germination, we planted seedlings in 6-cell seedling plug trays with soil mix (1:3 ratio of Promix soil and sand, 300mL nutricote fertilizer per 60 lbs), in a glasshouse at the University of Toronto. After 20 days, we transplanted individuals to 6-inch plastic pots which were watered every other day and fertilized with all-purpose plant food (Miracle-Gro) every 2 weeks. We selected healthy individuals for leaf tissue and plants were subjected to 24 hours in the dark prior to collection. We sampled young leaves which were flash froze in liquid nitrogen and stored at –80 °C before being sent to Cantata Bio (Scotts Valley, CA, US) for DNA extraction, library preparation and PacBio sequencing.

### Assembly and Annotation

We made use of our recently generated whole-genome phased assembly of an *R. hastatulus* XYY male, and the assembly of the hermaphroditic species *R. salicifolius* (Sacchi, Humphries et al. 2023). In addition to RNA-Seq, we generated a new long-read de novo genome assembly for the hermaphroditic species *Rumex bucephalaphorus* using a combination of high-coverage HiFi PAC Bio sequencing and Dovetail Omni-C sequencing. Finally, we generated transcriptome assemblies from RNASeq data for the other 9 species. To estimate genome size for coverage in *R. bucephalophorus* and 5 other species in our dataset, we performed flow cytometry conducted by Plant Cytometry Services, https://www.plantcytometry.nl/. Briefly, nuclei were stained with DAPI and DNA content per nucleus was quantified relative to *Vinca minor* as absolute DNA ratio with *Vinca minor* multiplied with DNA content of *Vinca minor* (1,51 pg/2C or 1477 Mbp/2C).

For the assembly of *R. bucephalophorus*, we grew field-collected seed in the University of Toronto glasshouse. 17.42 g of leaf tissue was used to extract high-molecular weight DNA by Dovetail Genomics (Cantata Bio, LLC, Scotts Valley, CA, USA). PAC Bio CCS reads (Pacific Biosciences Menlo Park, CA, USA) were sequenced by Dovetail for a total of ∼50 GB (approximately 26X coverage, based on a genome size estimate of 1.96 GB from flow cytometry (Supplementary Data 1). We used hifiasm-0.19.5 (Cheng et al. 2022) using the –primary assembly option to generate the contig-level assembly. Following this, paired-end OmniC reads were then mapped and filtered to the assembly using bwa v0.7.15 (Li and Durbin 2009) following the Arima mapping pipeline (https://github.com/ArimaGenomics/mapping_pipeline), and resulting filtered (MapQ>10) bam files had duplicates marked using Picard v2.7.1. We scaffolded the assembly using YaHS under default parameters (Zhou et al. 2023) to generate the final scaffolded assembly. Scaffolds 9 and 10 of our assembly, upon visual inspection of synteny with *R. salicifolius* and examination of synteny of the genome to itself (Supplementary Figures 1 and 2), are likely separately assembled heterozygous copies of other autosomes; these scaffolds were removed for downstream analyses, along with additional smaller scaffolds.

For the transcriptome assemblies, we used Trinity v2.11.0 (Grabherr et al. 2011) with each of the 10 species to create a transcriptome assembly for the sequencing libraries (representing a single tissue/individual combination). These assemblies were combined for each species and then reduced to representative species-wide transcriptomes using the traa2cds.pl script from EvidentialGene (Gilbert 2013) and retaining the primary transcript for each gene model.

### Orthogroup Identification and Alignment

From our final EviGene transcriptome assemblies, we used a custom shell script to filter out partial coding sequences (CDS) and CDSs belonging to classes other than the “main” class. For *R. hastatulus*, we used *gffread* (Pertea and Pertea 2020) to extract CDS from a previously published genome annotation (Rifkin et al. 2022). For use as an outgroup, we acquired previously published annotated coding sequences from the genome assembly of Tartary buckwheat, *Fagopyrum tataricum* (Zhang et al. 2017). This final set of CDSs for each species was used for downstream comparative analyses. To identify orthologous coding regions, we first used OrthoFinder v2.5.4 (Emms & Kelly 2019) with the default parameters to assign our assembled CDSs from each species into orthogroups. The sequences for each orthogroup were then translated to amino acids using BioPython (python v3.6.8) (Cock et al. 2009), and amino acid alignment was performed for each using MUSCLE v3.8.1551 (Edgar 2004). We used these amino acid alignments to guide codon alignment for each orthogroup using RevTrans 2.0 (Wernersson & Pedersen 2003), and quality filtering of the final codon alignments was conducted using Gblocks v0.91b (Castresana 2000) with the –b5 setting allowing for gap positions.

### Phylogenetic inference

Our OrthoFinder analysis identified very few (less than 100) single-copy orthologs across all 12 species in our dataset. Therefore, we employed two sampling schemes to increase the number of loci available for downstream phylogenetic inference. First, we allowed for some missing data, including orthogroups in which a copy was present in at least 9 of the 12 species. Second, we included orthogroups where gene duplication events were limited to a single species, as differential loss of duplicates cannot affect phylogenetic inference in such cases (Smith & Hahn 2021). In such orthogroups, we randomly sampled a single gene copy from the species containing duplicates for downstream inferences. This sampling scheme resulted in a dataset of 5,263 single-copy genes.

For maximum-likelihood inference, we generated a concatenated alignment of all 5,263 loci. This alignment was subsequently filtered using Gblocks for codon sequences, with the –b5 setting allowing for gap positions. The final filtered alignment contained 6.5 Mb of coding sequence. This alignment was given to IQ-TREE v2.1.2 to infer a phylogeny using ModelFinder, SH-aLRT, and ultrafast bootstrap with 1000 replicates. ModelFinder uses maximum-likelihood inference to estimate the best-fitting model of sequence evolution for the data, while SH-aLRT and ultrafast bootstrap are alternative methods for assessing branch support.

Biological sources of gene tree discordance – incomplete lineage sorting and introgression – can mislead maximum-likelihood approaches to phylogenetic inference (Degnan and Rosenberg 2006, Mendes and Hahn 2018). To quantify the degree of gene tree discordance in our dataset, and its potential effects on our inferred phylogeny, we used the gene trees inferred in the previous section for our set of single-copy orthogroups. We calculated both gene (gCF) and site concordance factors (sCF) on our maximum-likelihood tree using functions available in IQ-TREE (Minh et al. 2020b, Mo et al. 2023). These measure the proportion of gene trees and parsimony-informative sites, respectively, that support a particular branch in the inferred species tree. We also inferred a phylogeny using ASTRAL-III (Zhang et al. 2018), a summary approach that is robust to the effects of incomplete lineage sorting.

We time-calibrated our maximum-likelihood phylogeny using a penalized likelihood approach implemented in the *chronos* function of the R package *ape* (Sanderson 2002). The split of the XYY clade and the *R. bucephalophorus*/XY clade was constrained to 10.8 MYA, and the split of the ancestor of those two clades from the clade containing *R. sagittatus* and *R. scutatus* was constrained to 13.77 MYA, based on date estimates from a recent study (Koenemann et al. 2023). We constrained the root of the *Rumex* clade to a maximum age of 23 MYA following Koenemann et al. 2023, based on fossil evidence (Muller 1981, Barrón et al., 2006, Huang et al. 2022). We fit correlated, discrete, and relaxed clock models, and the model producing the lowest PHIIC score was chosen as the best-fitting.

### Introgression analysis

Post-speciation introgression is expected to cause an asymmetry in discordant gene tree frequencies, forming the basis for many common tests, such as the *D*-statistic (Green et al. 2010, Durand et al. 2011). To investigate the prevalence of introgression both before and after the origins of sex chromosomes, we used the Δ test, which tests for an asymmetry in gene tree counts directly, and allows ancestral branches of the phylogeny to be tested (Huson et al. 2005, Vanderpool et al. 2020). Given a rooted triplet where the species phylogeny is [(A,B),C], the two discordant topologies are [(B,C),A] and [(A,C),B]. A, B, and C can be monophyletic clades (when testing ancestral branches) or individual species (when testing introgression among contemporary species). The Δ test is then simply the following:

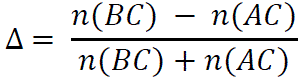

or, the difference in count between the two discordant topologies, divided by their sum. The statistic is normalized between –1 and 1, with the sign indicating the branches involved in introgression.

In applying the test to *Rumex*, we made use of our set of gene trees inferred from single-copy orthogroups. We tested all possible rooted triplets in the phylogeny, using *F. tataricum* to root each gene tree. In cases where a set of significant tests between multiple species pairs could be explained by introgression between their common ancestral populations, we favored the ancestral event on the basis of parsimony. We evaluated significance with bootstrapping: for each test, we randomly sampled 1000 datasets of 5,263 gene trees each from our original dataset with replacement. Δ was recalculated for each bootstrapped dataset to generate a sampling distribution, and significance was assessed by calculating the degree to which the distribution overlapped 0 (the null hypothesis). For a mean of the sampling distribution > 0, this would be the proportion of Δ values in the distribution < 0, and vice-versa. Tests with a very small number of discordant gene trees (less than 5% of the total) were discarded.

In addition to the Δ statistic, we estimated phylogenetic networks using the software *PhyloNet* v3.8.2 (Than et al. 2008). A phylogenetic network is a tree structure that includes horizontal reticulation edges connecting lineages, which are used to represent introgression events. Computational limitations prevented us from using the full-likelihood inference, so we applied the topology-based pseudolikelihood method *InferNetwork_MPL* (Yu & Nakhleh 2015) to our dataset of 5,263 gene tree topologies, using the default parameter settings and allowing a maximum of 6 reticulations.

### Resolving the history of sex chromosome evolution

To assess X chromosome homology between the XY and XYY clades, we leveraged a previously generated list of sex-linked genes in *R. rothschildianus* using SNP segregation patterns from transcriptome data (Crowson et al. 2017), as well as our genome assembly for XYY *R. hastatulus* (Sacchi, Humphries et al. 2023). We used BLASTn (Altschul et al. 1990) against the maternal (X-bearing) haplotype and used the top scoring BLAST hit to identify the location of both X-hemizygous genes (genes with the Y either silenced or deleted) and those with a Y gametolog from *R. rothschildianus*. We plotted the density of BLAST hits against the genome of *R. hastatulus* in 25 evenly spaced windows along each chromosome.

We investigated the plausibility of loss of XY sex chromosomes in *R. bucephalophorus* by examining the relationship of *R. bucephalophorus* to *R. hastatulus* X/Y gametologs. First, we used custom Python scripts to extract X and Y-linked coding sequences from our phased and annotated XYY *R. hastatulus* genome assembly, all coding sequences from our genome assemblies of *R. bucephalophorus* and *R. salicifolius*. and transcriptome assemblies of *R. trisetifer* and *R. amurensis*. We ran OrthoFinder on these six datasets to identify orthogroups, and extracted usable single-copy orthogroups where duplications were limited to one sample. Gene trees were rooted on one of *R. salicifolius*, *R. trisetifer*, or *R. amurensis*, depending on presence/absence of data; in trees where multiple of these species were present, one was chosen at random to root the tree. These orthogroups were subsequently aligned with MUSCLE and gene trees were estimated using IQ-TREE. We then used the *ete3* package (Huerta-Cepas et al. 2016) implemented in Python to tally the three possible gene tree topologies: no loss (X and Y gametologs sister), loss of Y (*R. bucephalophorus* sister to X), and loss of X (*R. bucephalophorus* sister to Y).

## Supplementary Figures

**Supplementary Figure 1:**
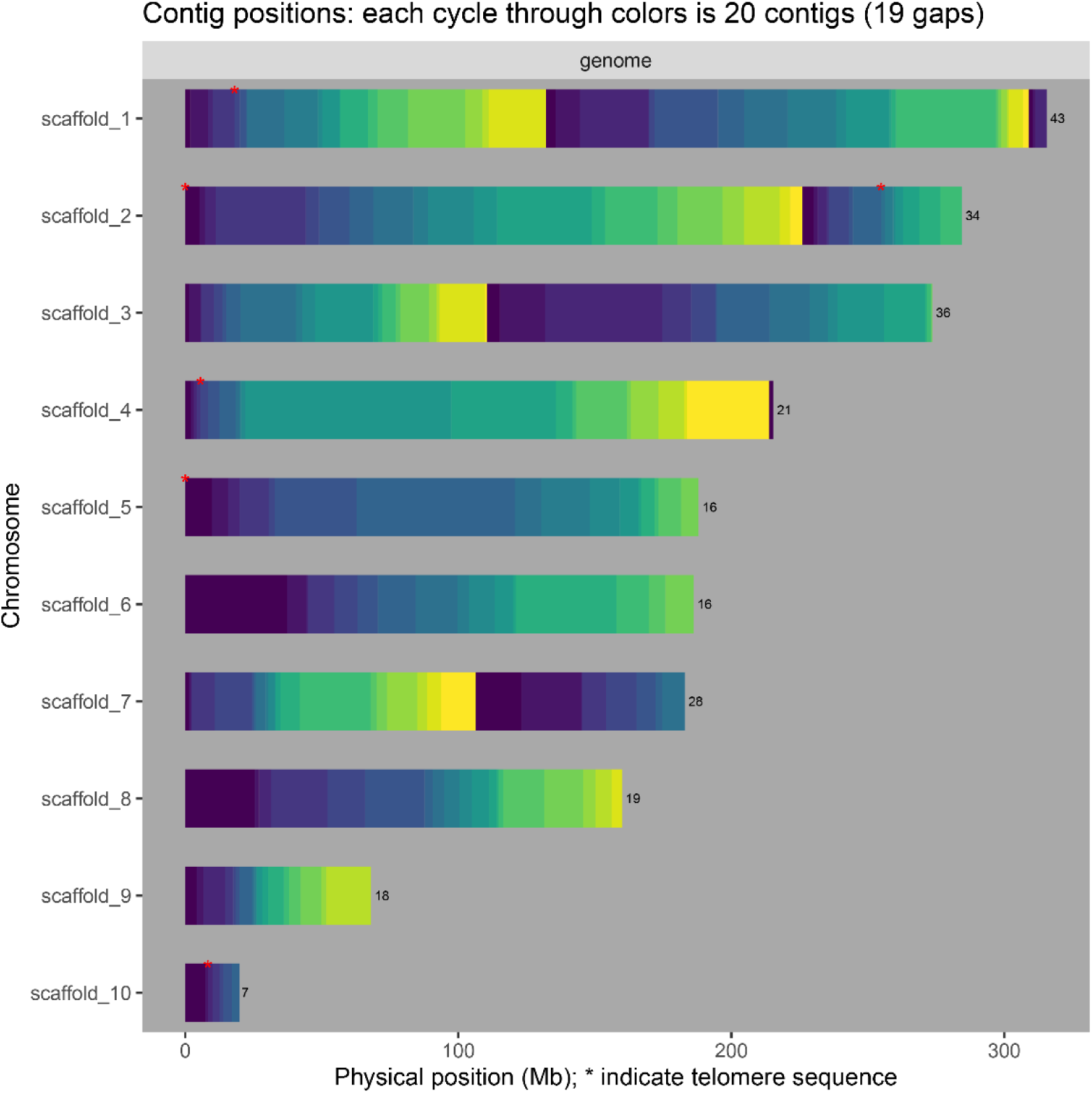
Genome assembly summary for *R. bucephalophorus*, with scaffolds order by physical size. Each color cycle (from dark blue to yellow) represents 20 contigs. Scaffolds 9 and 10 likely represent misassembled heterozygous copies of other chromosomes (see Supplementary Figures 2 and 3); karyotyping indicates *R. bucephalophorus* has 8 chromosomes (Navajas-Pérez et al. 2005).

**Supplementary Figure 2:**
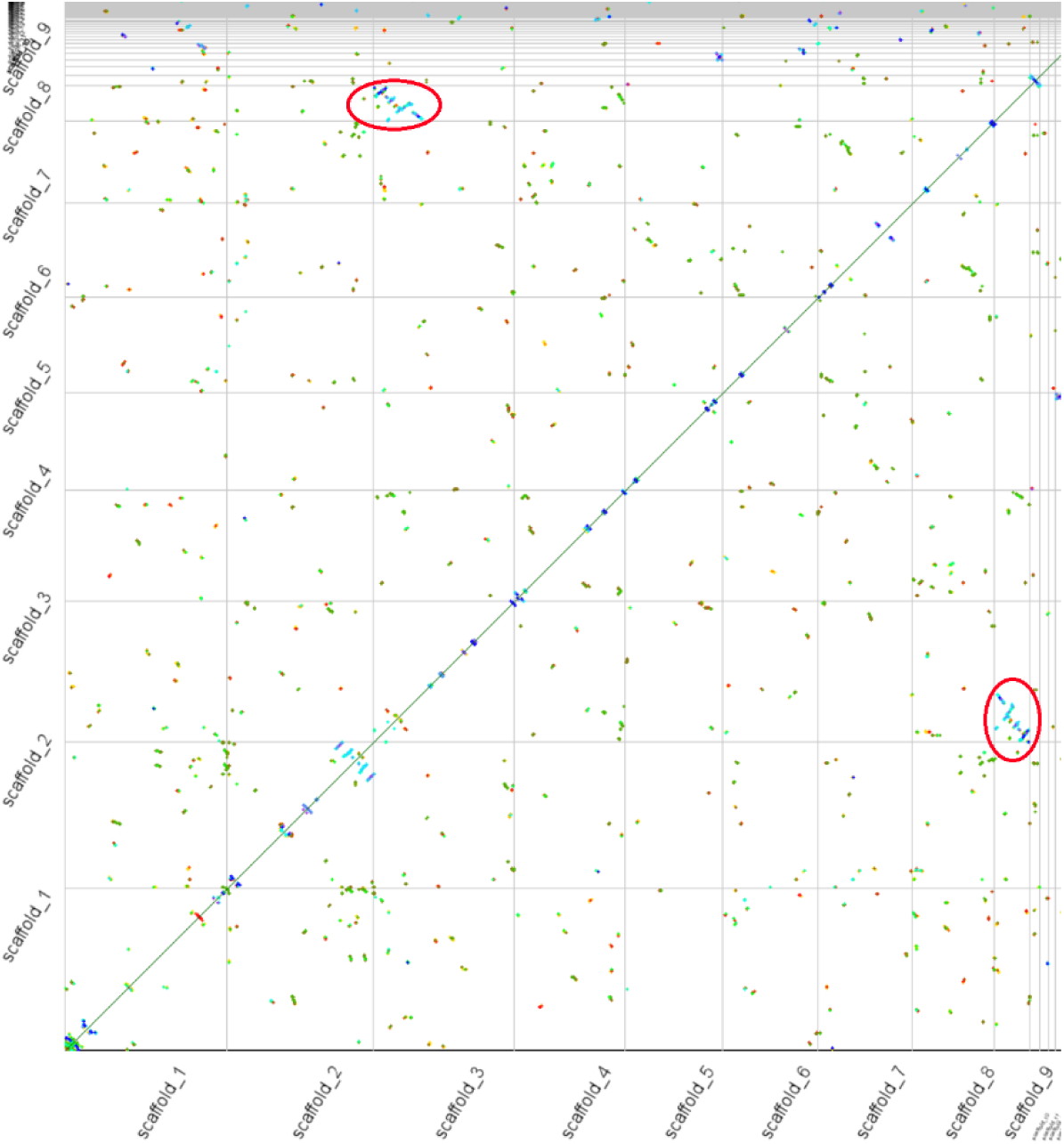
Synteny dotplot of our assembly of *R. bucephalophorus* against itself. Scaffold 9 shows high similarity to scaffold 3 (red circles), indicating a separately assembled heterozygous copy of scaffold 3.

**Supplementary Figure 3:**
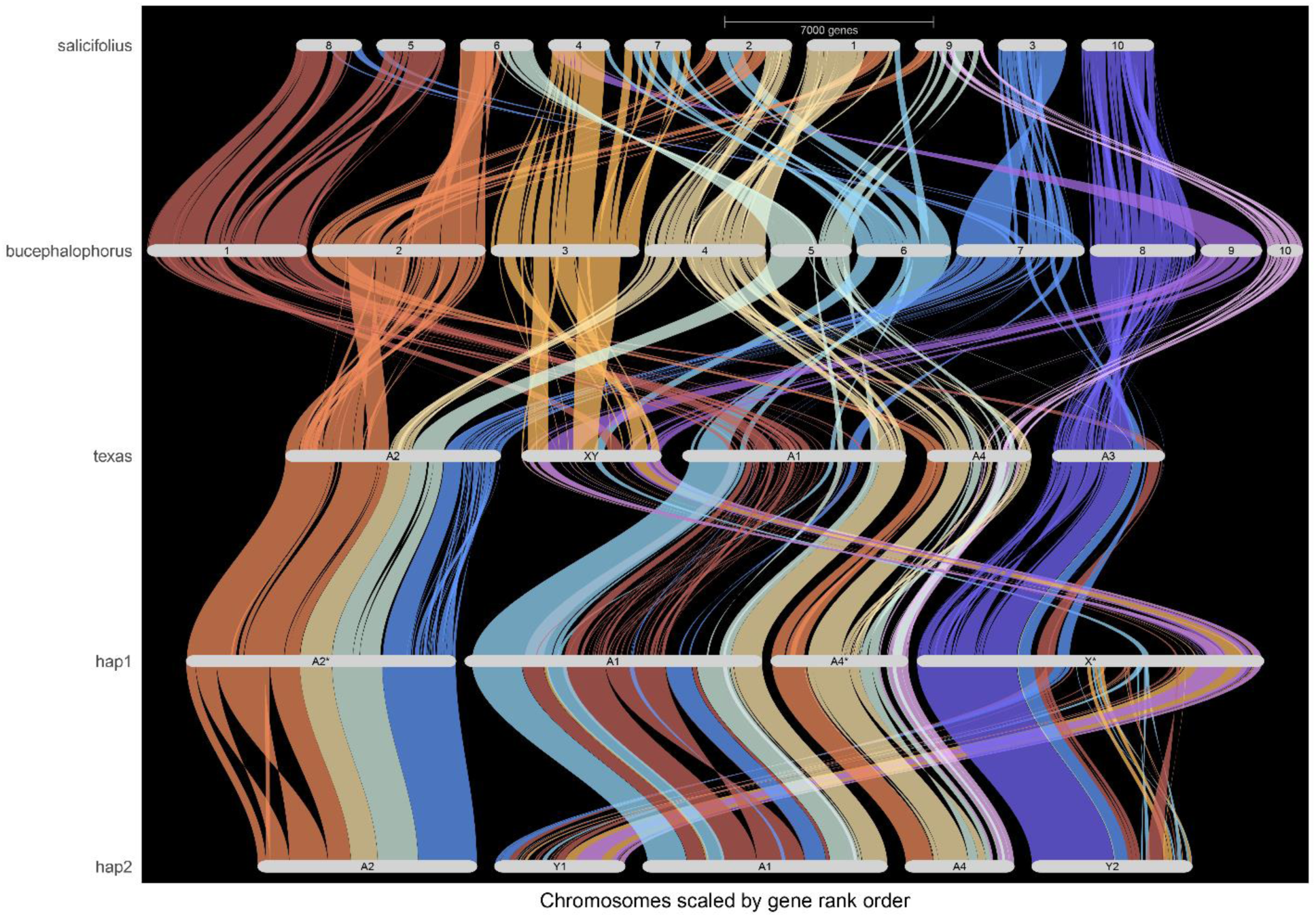
GENESPACE synteny plot of *R. salicifolius*, *R. bucephalophorus*, and the two cytotypes of *R. hastatulus*, including the extra assembled scaffolds 9 and 10 of *R. bucephalophorus*. These scaffolds are syntenic to regions of *R. salicifolius* that overlap with other chromosomes, again indicating extra assembled copies of these chromosomes.

**Supplementary Figure 4:**
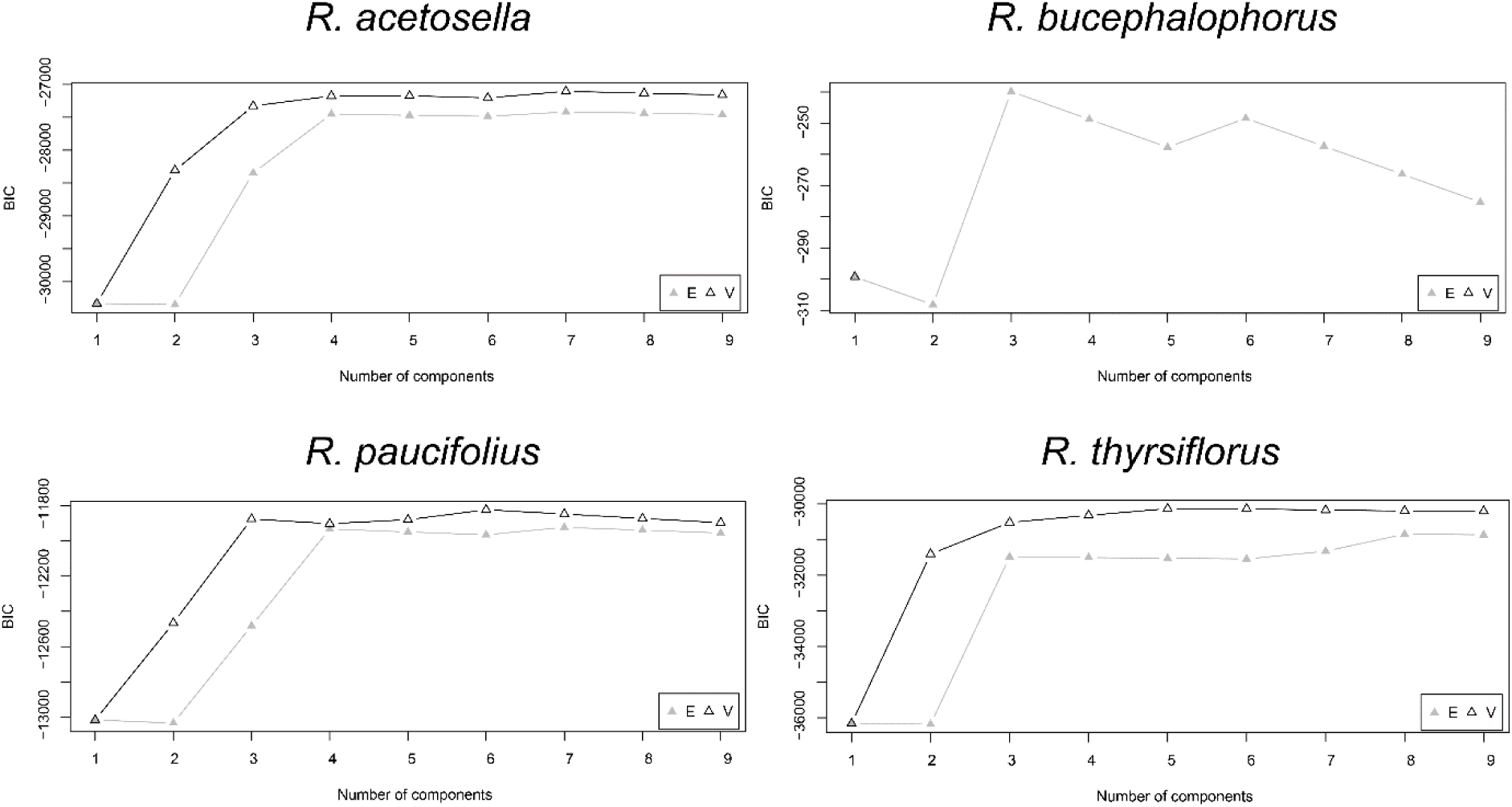
BIC values by number of components fit to the distribution of log(dS) values, estimated by *mclust*. For clarity, results only shown for species where support for 2 components was found (*R. acetosella/bucephalophorus/paucifolius/thyrsiflorus*). In each plot, the “E” and “V” lines denote BIC scores for equal-variance and unequal-variance models for each # of components.

**Supplementary Figure 5:**
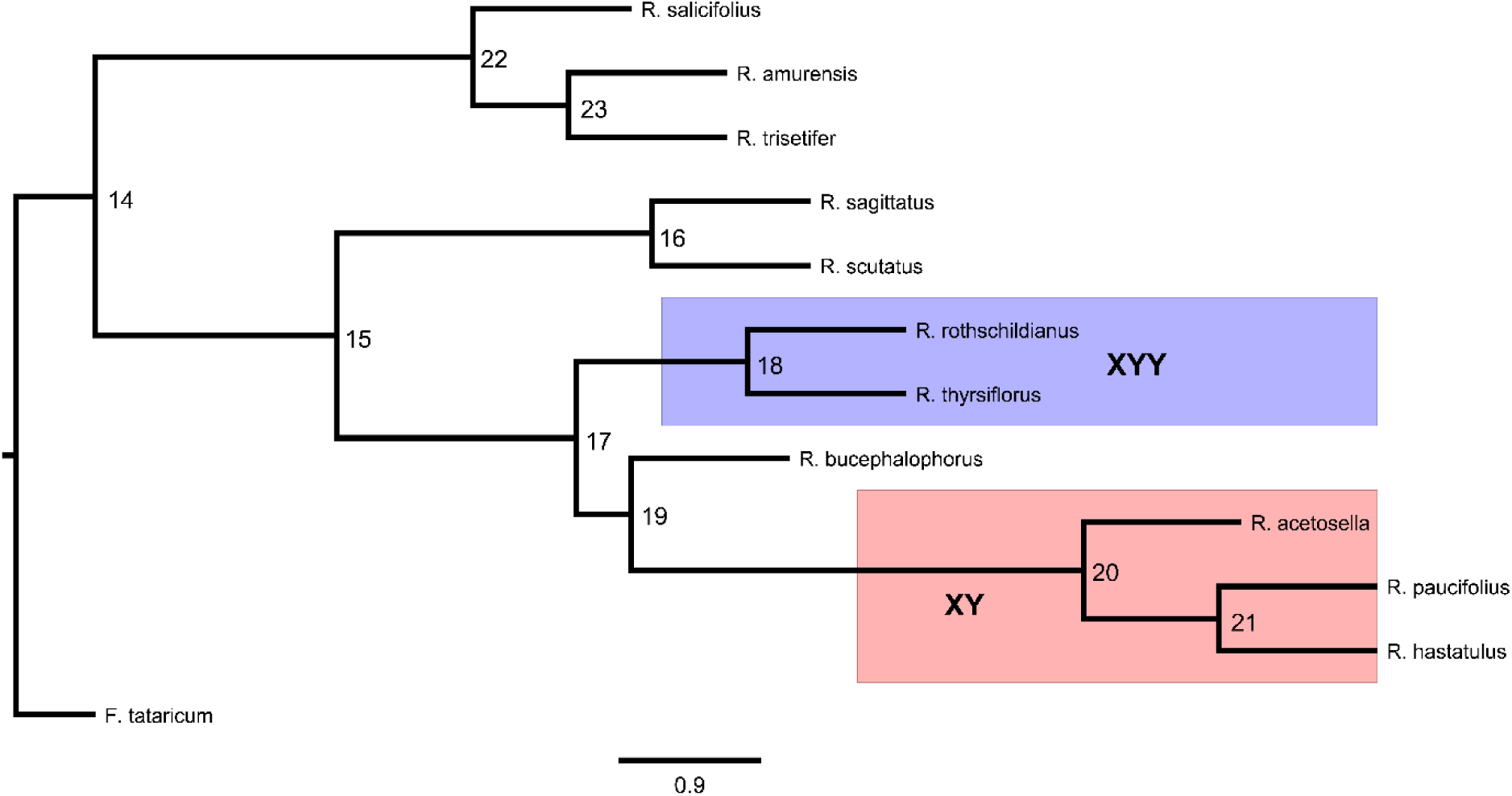
ASTRAL-III species tree topology for *Rumex* inferred from 5,263 gene orthologs. Nodes are labelled with their traversal order (for interpreting Supplementary Tables 1 and 2).

**Supplementary Figure 6:**
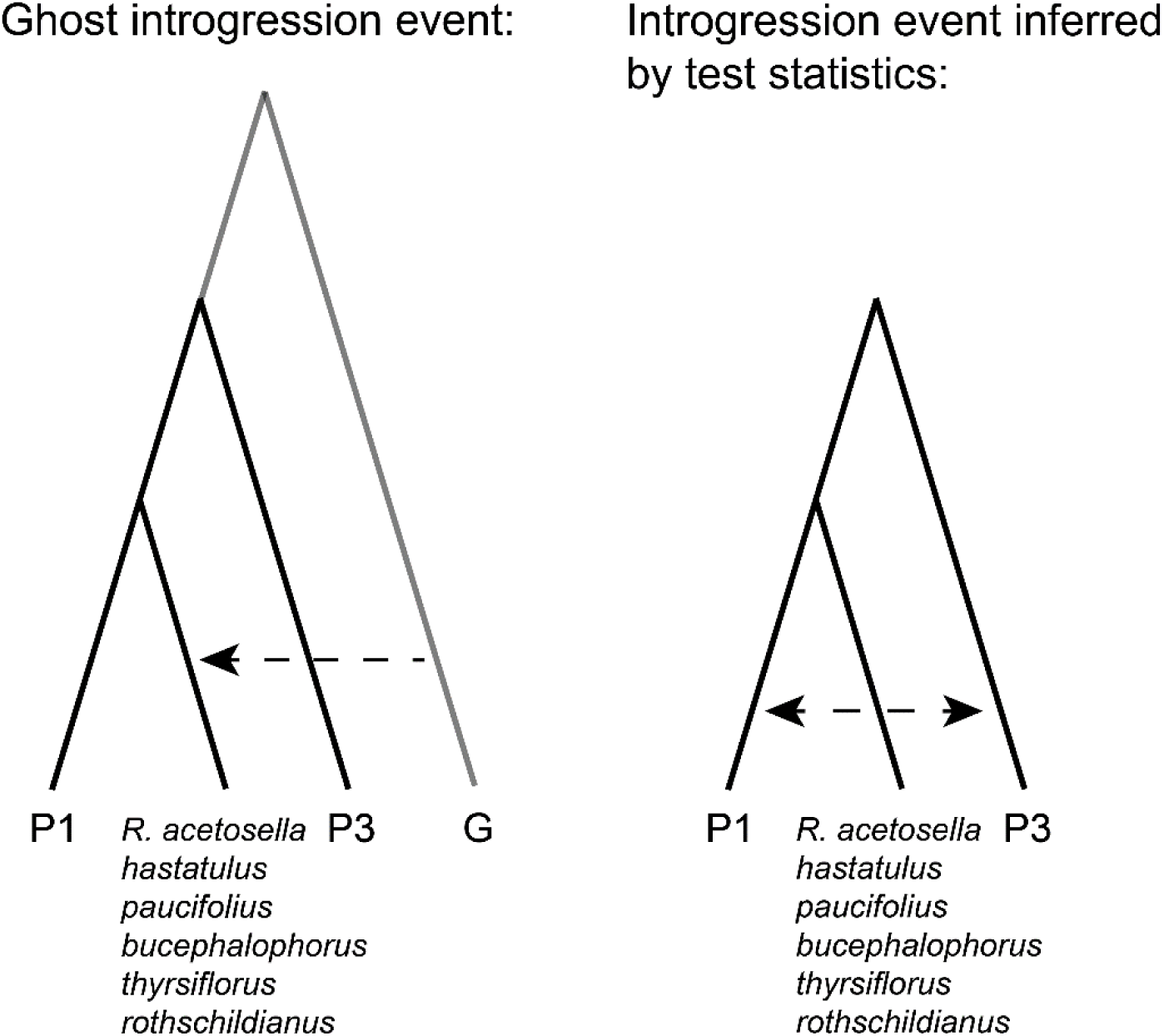
Inferring ghost introgression from test statistics based on gene tree topologies. In a scenario where a diverged unsampled population introgresses into one of the indicated species (lefthand side), introgression test statistics will imply introgression between the other two species used in the test (i.e. P1 and P3; righthand side). This happens because the presence of diverged alleles in the recipient species makes it appear less closely related to P1 than expected based on phylogenetic relationships. Importantly, this pattern occurs regardless of the identity of P1 and P3, distinguishing it from instances of true introgression between any particular combination of P1 and P3. We observe this pattern for the species indicated in the figure (see Supplementary Table X).

**Supplementary Figure 7:**
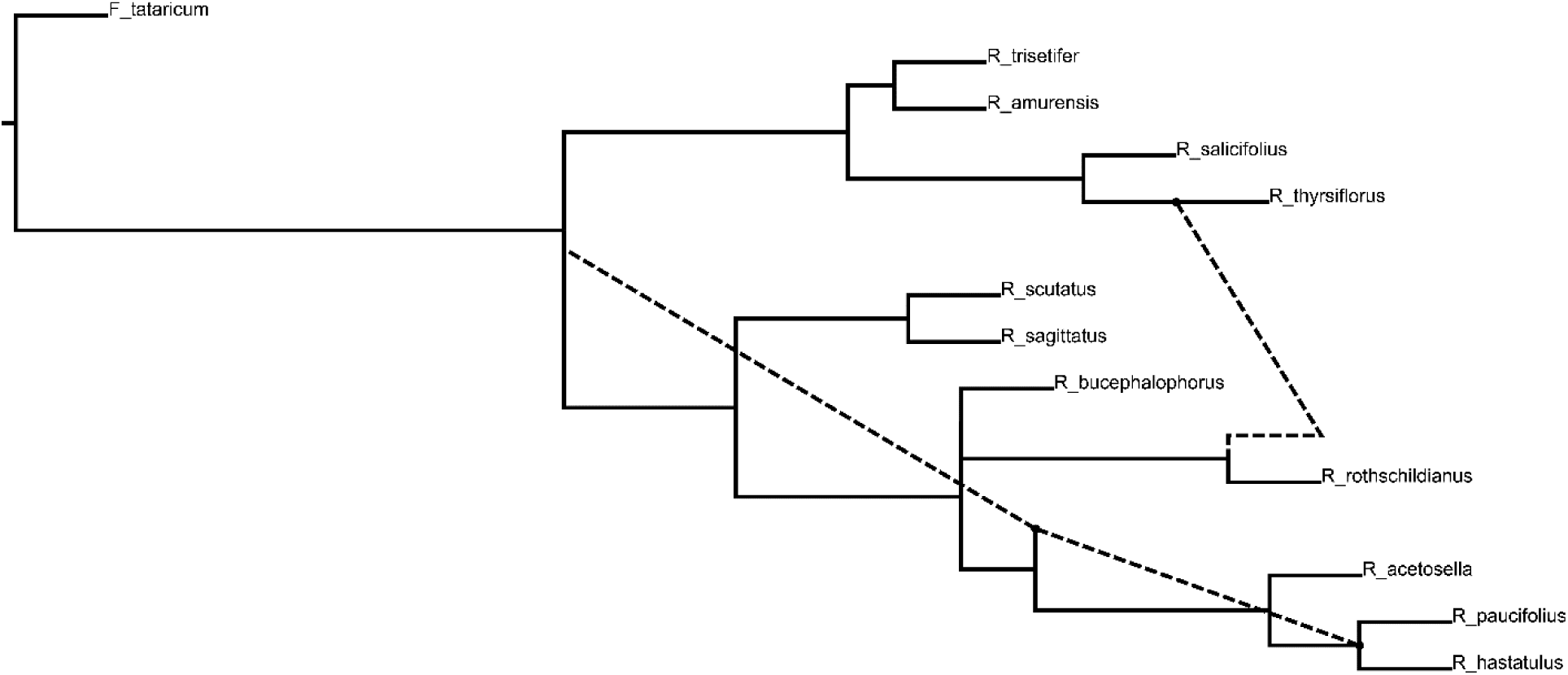
Best-fitting phylogenetic network for *Rumex* inferred by PhyloNet’s pseudolikelihood method. Proposed reticulations are indicated by the dashed lines.

## Supplementary Tables

**Supplementary Table 1:**
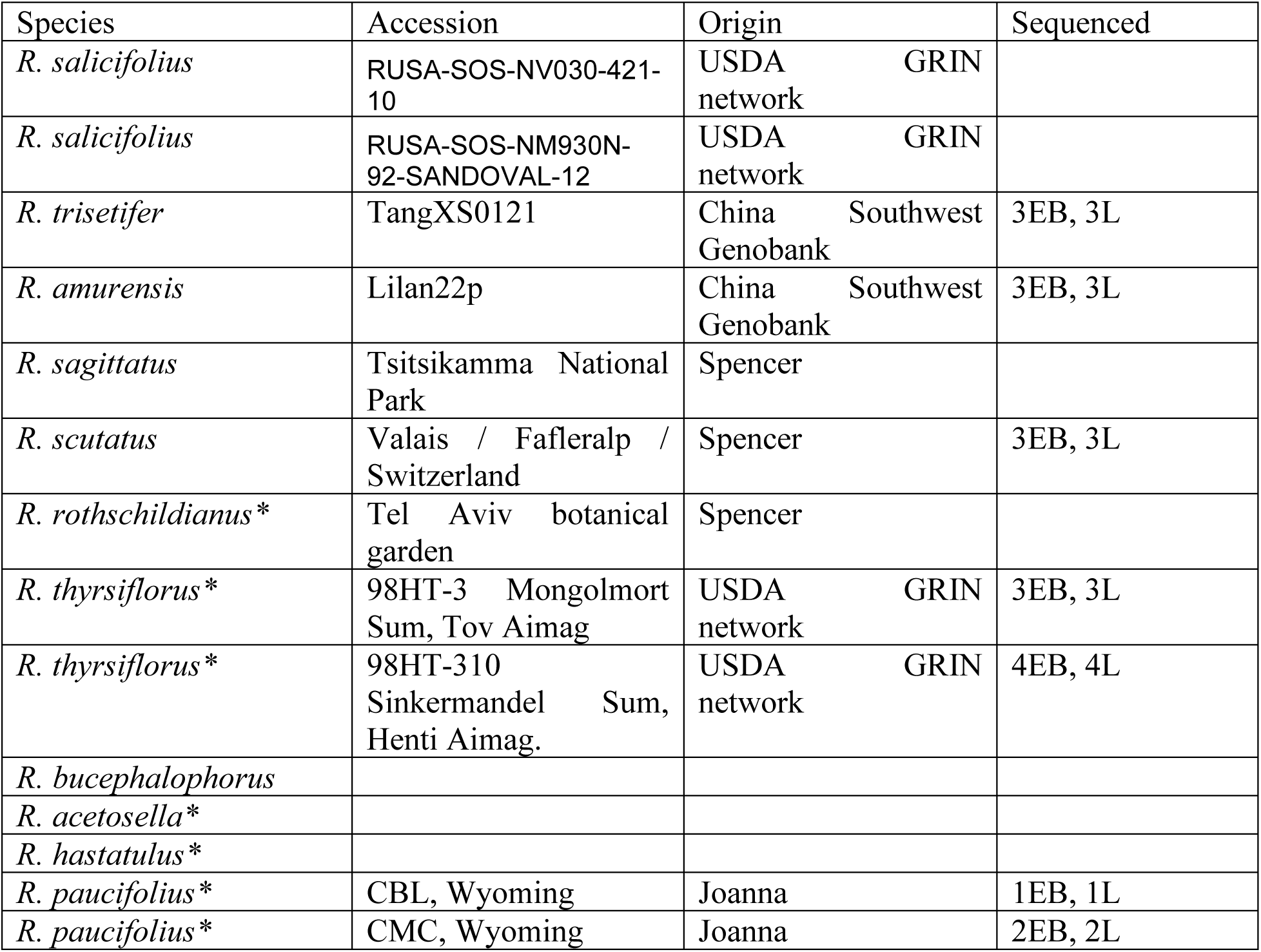
Sampling origins of sequenced accessions. Under the “sequenced” column, “P” indicates pollen, “EB” indicates early buds, and “L” indicates leaves. Numbers correspond to individuals sampled for each tissue. Dioecious species are indicated with an asterisk.

**Supplementary Table 2:** Genome assembly summary statistics for *R. bucephalophorus*.

|  |  |
| --- | --- |
| **Contig L50** | 45 |
| **Contig N50** | 13660000 |
| **Scaffold L50** | 3 |
| **Scaffold N50** | 215375990 |

**Supplementary Table 3:** Gene concordance factors for each node in the *Rumex* phylogeny (nodes labelled in Supplementary Figure 5). gDF1 and gDF2 indicate the frequencies for the most common and second most common discordant gene tree topologies, respectively. gN indicates the number of gene trees used to calculate each value.

| **ID** | **gCF** | **gDF1** | **gDF2** | **gN** |
| --- | --- | --- | --- | --- |
| 14 | 100 | 0 | 0 | 5261 |
| 15 | 68.18 | 4.19 | 3.25 | 5255 |
| 16 | 88.86 | 2.09 | 4.09 | 4695 |
| 17 | 72.82 | 0.55 | 2.28 | 6005 |
| 18 | 76.09 | 19.72 | 1.12 | 5620 |
| 19 | 47.08 | 21.9 | 17.02 | 1680 |
| 20 | 93.69 | 0.88 | 1.33 | 1584 |
| 21 | 69.97 | 15.07 | 12.99 | 1878 |
| 22 | 74.66 | 0.72 | 1.2 | 5011 |
| 23 | 61.34 | 26.25 | 1.57 | 4446 |

**Supplementary Table 4:** Site concordance factors for each node in the *Rumex* phylogeny (nodes labelled in Supplementary Figure 5). sDF1 and sDF2 indicate the frequencies for the most common and second most common discordant site patterns, respectively. sN indicates the mean number of informative sites used to calculate each value.

| **ID** | **sCF** | **sDF1** | **sDF2** | **sN** |
| --- | --- | --- | --- | --- |
| 15 | 47.02 | 27.13 | 25.85 | 118404.5 |
| 16 | 62.98 | 18.02 | 19.01 | 114758.3 |
| 17 | 56.46 | 17.42 | 26.12 | 103928.7 |
| 18 | 60 | 31.12 | 8.88 | 110316.9 |
| 19 | 45.19 | 28.43 | 26.37 | 27991.59 |
| 20 | 83.65 | 8.76 | 7.59 | 28369.21 |
| 21 | 53.73 | 14.32 | 31.95 | 16367.81 |
| 22 | 84.4 | 8.35 | 7.25 | 112043 |
| 23 | 56.58 | 37.1 | 6.32 | 46551.48 |

## Descriptions of Supplementary Data Files

*Supplementary Data 1*: Flow cytometry results for R. *acetosella*, R. *paucifolius*, R. *bucephalophorus*, R. *thyrsiflorus*, R. *sagittatus*, R. *salicifolius*, and several unsequenced species.

*Supplementary Data 2*: Transcriptome assembly summary statistics for our ten sequenced study species.

*Supplementary Data 3*: Δ statistic results applied to *Rumex* transcriptomic data. The P1, P2, and P3 columns indicate the two sister species and the unpaired species in the test, respectively. All tests used tartary buckwheat (*F. tataricum*) as outgroup. A positive test implies introgression between P1 and P3, while a negative test implies introgression between P2 and P3; however, the interpretation of many of these tests is complicated by our proposed ghost introgression events (Figure 2, Supplementary Figures 4 and 5, section “*Signatures of ghost introgression in the Rumex phylogeny*” in the main text).

## References

1. Adhikari, K. N., Campbell, C. G. (1998). In vitro germination and viability of buckwheat (*Fagopyrum esculentum* Moench) pollen. Euphytica, 102, 87–92. doi:10.1023/A:1018393425407

2. Altschul, S. F., Gish, W., Miller, W., Myers, E. W., & Lipman, D. J. (1990). Basic local alignment search tool. Journal of Molecular Biology, 215(3), 403–410. doi:10.1016/S0022-2836(05)80360-2

3. Anderson, N. W., Hjelmen, C.E., Blackmon, H. (2020). The probability of fusions joining sex chromosomes and autosomes. Biology Letters, 16(11), 20200648. doi:10.1098/rsbl.2020.0648

4. Avise, J. C., Robinson, T. J. (2008). Hemiplasy: a new term in the lexicon of phylogenetics. Systematic Biology, 57(3): 503–507. doi:10.1080/10635150802164587

5. Bachtrog, D. (2013). Y-chromosome evolution: emerging insights into processes of Y-chromosome degeneration. Nature Reviews Genetics, 14(2), 113–124. doi:10.1038/nrg3366

6. Bachtrog, D., Mank, J. E., Peichel, C. L., Kirkpatrick, M., Otto, S. P., Ashman, T. L., … Consortium, T. S. (2014). Sex determination: why so many ways of doing it? PLoS Biology, 12(7). doi:10.1371/journal.pbio.1001899

7. Barrett, S.C.H. (2002). The evolution of plant sexual diversity. Nature Reviews Genetics, 3, 274–284. doi:10.1038/nrg776

8. Barron, E., Lassaletta, L., & Alcalde-Olivares, C. (2006). Changes in the Early Miocene palynoflora and vegetation in the east of the Rubielos de Mora Basin (SE Iberian Ranges, Spain). Neues Jahrbuch Fur Geologie Und Palaontologie-Abhandlungen, 242(2-3), 171–204. doi:10.1127/njgpa/242/2006/171

9. Beaudry, F.E.G., Barrett., S.C.H., & Wright, S.I. (2020). Ancestral and neo-sex chromosomes contribute to population divergence in a dioecious plant. Evolution, 74(2): 256–269. doi:10.1111/evo.13892

10. Beaudry, F. E. G., Rifkin, J. L., Peake, A. L., Kim, D., Jarvis-Cross, M., Barrett, S. C. H., & Wright, S. I. (2022). Effects of the neo-X chromosome on genomic signatures of hybridization in *Rumex hastatulus*. Molecular Ecology, 31(13), 3708–3721. doi:10.1111/mec.16496

11. Bergero, R., Forrest, A., Kamau, E., & Charlesworth, D. (2007). Evolutionary strata on the X chromosomes of the dioecious plant Silene latifolia: evidence from new sex-linked genes. Genetics, 175(4), 1945–1954. doi:10.1534/genetics.106.070110

12. Bergero, R., Gardner, J., Bader, B., Yong, L., & Charlesworth, D. (2019). Exaggerated heterochiasmy in a fish with sex-linked male coloration polymorphisms. Proceedings of the National Academy of Sciences of the United States of America, 116(14), 6924–6931. doi:10.1073/pnas.1818486116

13. Blanc, G., & Wolfe, K. H. (2004). Widespread paleopolyploidy in model plant species inferred from age distributions of duplicate genes. Plant Cell, 16(7), 1667–1678. doi:10.1105/tpc.021345

14. Bracewell, R. R., Bentz, B. J., Sullivan, B. T., & Good, J. M. (2017). Rapid neo-sex chromosome evolution and incipient speciation in a major forest pest. Nature Communications, 8(1), 1593. doi:10.1038/s41467-017-01761-4

15. Buchfink, B., Reuter, K., & Drost, H. G. (2021). Sensitive protein alignments at tree-of-life scale using DIAMOND. Nature Methods 18: 366–368. doi: 10.1038/s41592-021-01101-x

16. Bull, J. J. (1983). Evolution of sex determining mechanisms. Menlo Park, CA: Benjamin Cummings.

17. Bull, J. J., & Charnov, E. L. (1977). Changes in the heterogametic mechanism of sex determination. Heredity, 39(1), 1–14. doi:10.1038/hdy.1977.38

18. Castillo, E. R. D., Taffarel, A., & Marti, D. A. (2014). The early evolutionary history of neo-sex chromosomes in Neotropical grasshoppers, *Boliviacris noroestensis* (Orthoptera: Acrididae: Melanoplinae). European Journal of Entomology, 111(3), 321–327. doi:DOI 10.14411/eje.2014.047

19. Castresana, J. (2000). Selection of conserved blocks from multiple alignments for their use in phylogenetic analysis. Molecular Biology and Evolution, 17(4), 540–552. doi:10.1093/oxfordjournals.molbev.a026334

20. Charlesworth, B. (1991). The evolution of sex chromosomes. Science, 251(4997), 1030–1033. doi:10.1126/science.1998119

21. Charlesworth, B., & Charlesworth, D. (1978). A model for the evolution of dioecy and gynodioecy. The American Naturalist, 112(988), 975–997. doi:10.1086/283342

22. Charlesworth, B., & Charlesworth, D. (2000). The degeneration of Y chromosomes. Philosophical Transactions of the Royal Society of London B: Biological Sciences, 355(1403), 1563–1572. doi:10.1098/rstb.2000.0717

23. Charlesworth, B., Coyne, J. A., & Barton, N. H. (1987). The relative rates of evolution of sex-chromosomes and autosomes. American Naturalist, 130(1), 113–146. doi:10.1086/284701

24. Charlesworth, D. (2013). Plant sex chromosome evolution. Journal of Experimental Botany, 64(2), 405–420. doi:10.1093/jxb/ers322

25. Charlesworth, D., & Charlesworth, B. (1980). Sex differences in fitness and selection for centric fusions between sex-chromosomes and autosomes. Genetics Research, 35(2), 205–214. doi:10.1017/s0016672300014051

26. Charlesworth, D., Charlesworth, B., & Marais, G. (2005). Steps in the evolution of heteromorphic sex chromosomes. Heredity, 95(2), 118–128. doi:10.1038/sj.hdy.6800697

27. Cheng, H., Jarvis, E. D., Fedrigo, O., Koepfli, K. P., Urban, L., Gemmell, N. J., Li, H. (2022). Haplotype-resolved assembly of diploid genomes without parental data. Nature Biotechnology 40, 1332–1335. doi: 10.1038/s41587-022-01261-x

28. Cock, P. J., Antao, T., Chang, J. T., Chapman, B. A., Cox, C. J., Dalke, A., … de Hoon, M. J. (2009). Biopython: freely available Python tools for computational molecular biology and bioinformatics. Bioinformatics, 25(11), 1422–1423. doi:10.1093/bioinformatics/btp163

29. Connallon, T., Olito, C., Dutoit, L., Papoli, H., Ruzicka, F., & Yong, L. (2018). Local adaptation and the evolution of inversions on sex chromosomes and autosomes. Philosophical Transactions of the Royal Society of London B: Biological Sciences, 373(1757). doi:10.1098/rstb.2017.0423

30. Crowson, D., Barrett, S. C. H., & Wright, S. I. (2017). Purifying and positive selection influence patterns of gene loss and gene expression in the evolution of a plant sex chromosome system. Molecular Biology and Evolution, 34(5), 1140–1154. doi:10.1093/molbev/msx064

31. Degnan, J. H., & Rosenberg, N. A. (2006). Discordance of species trees with their most likely gene trees. PLoS Genetics, 2(5), e68. doi:10.1371/journal.pgen.0020068

32. Degnan, J. H., & Rosenberg, N. A. (2009). Gene tree discordance, phylogenetic inference and the multispecies coalescent. Trends in Ecology and Evolution, 24(6), 332–340. doi:10.1016/j.tree.2009.01.009

33. Durand, E. Y., Patterson, N., Reich, D., & Slatkin, M. (2011). Testing for ancient admixture between closely related populations. Molecular Biology and Evolution, 28(8), 2239–2252. doi:10.1093/molbev/msr048

34. Edgar, R. C. (2004). MUSCLE: multiple sequence alignment with high accuracy and high throughput. Nucleic Acids Research, 32(5), 1792–1797. doi:10.1093/nar/gkh340

35. El Taher, A., Ronco, F., Matschiner M., Salzburger, W., & Bohne, A. (2021). Dynamics of sex chromosome evolution in a rapid radiation of cichlid fishes. Science Advances, 7(36), eabe8215. doi:10.1126/sciadv.abe8215

36. Emms, D. M., & Kelly, S. (2019). OrthoFinder: phylogenetic orthology inference for comparative genomics. Genome Biology, 20(1), 238. doi:10.1186/s13059-019-1832-y

37. Fawcett, J. A., Takeshima, R., Kikuchi, S., Yazaki, E., Katsube-Tanaka, T., Dong, Y., … Yasui, Y. (2023). Genome sequencing reveals the genetic architecture of heterostyly and domestication history of common buckwheat. Nature Plants, 9, 1236–1251. doi: 10.1038/s41477-023-01474-1

38. Fridolfsson, A. K., Cheng, H., Copeland, N. G., Jenkins, N. A., Liu, H. C., Raudsepp, T., … Ellegren, H. (1998). Evolution of the avian sex chromosomes from an ancestral pair of autosomes. Proceedings of the National Academy of Sciences of the United States of America, 95(14), 8147–8152. doi:10.1073/pnas.95.14.8147

39. Gilbert, D. (2013). Gene-omes built from mRNA seq not genome DNA. 7th annual Arthropod Genomics Symposium. Notre Dame. doi:10.7490/f1000research.1112594.1

40. Grabowska-Joachimiak, A., Kula, A., Ksiazczyk, T., Chojnicka, J., Sliwinska, E., & Joachimiak, A. J. (2015). Chromosome landmarks and autosome-sex chromosome translocations in *Rumex hastatulus*, a plant with XX/XY_1_Y_2_ sex chromosome system. Chromosome Research, 23(2), 187–197. doi:10.1007/s10577-014-9446-4

41. Grabherr, M. G., Haas, B. J., Yassour, M., Levin, J. Z., Thompson, D. A., Amit, I., … Regev, A. (2011). Trinity: reconstructing a full-length transcriptome without a genome from RNA-Seq data. Nature Biotechnology 23(7): 644–652.

42. Grant, K. D., Koenemann, D., Mansaray, J., Ahmed, A., Khamar, H., El Oualidi, J., & Burke, J. M. (2022). A new phylogeny of *Rumex* (Polygonaceae) adds evolutionary context to the diversity of reproductive systems present in the genus. PhytoKeys, 204, 57–72. doi:10.3897/phytokeys.204.85256

43. Graves, J. A., & Watson, J. M. (1991). Mammalian sex chromosomes: evolution of organization and function. Chromosoma, 101(2), 63–68. doi:10.1007/BF00357055

44. Green, R. E., Krause, J., Briggs, A. W., Maricic, T., Stenzel, U., Kircher, M., … Paabo, S. (2010). A draft sequence of the Neandertal genome. Science, 328(5979), 710–722. doi:10.1126/science.1188021

45. Guerrero, R. F., & Kirkpatrick, M. (2014). Local adaptation and the evolution of chromosome fusions. Evolution 68(10), 2747–2756. doi:10.1111/evo.12481

46. Guo, L., Bloom, J. S., Dols-Serrate, D., Boocock, J., Ben-David, E., Schubert, O. T., … Kruglyak, L. (2022). Island-specific evolution of a sex-primed autosome in a sexual planarian. Nature, 606(7913), 329–334. doi:10.1038/s41586-022-04757-3

47. Handley, L. J., Ceplitis, H., & Ellegren, H. (2004). Evolutionary strata on the chicken Z chromosome: implications for sex chromosome evolution. Genetics, 167(1), 367–376. doi:10.1534/genetics.167.1.367

48. Hough, J., Hollister, J.D., Wang, W., Barrett, S.C.H., Wright, S.I. (2014). Genetic degeneration of old and young Y chromosomes in the flowering plant *Rumex hastatulus*. Proceedings of the National Academy of Sciences of the United States of America 111(21): 7713–7718. doi:10.1073/pnas.1319227111

49. Huang, Y. J., Zhu, H., Su, T., Spicer, R. A., Hu, J. J., Jia, L. B., & Zhou, Z. K. (2022). Rise of herbaceous diversity at the southeastern margin of the Tibetan Plateau: first insight from fossils. Journal of Systematics and Evolution, 60(5), 1109–1123. doi:10.1111/jse.12755

50. Huerta-Cepas, J., Serra, F., & Bork, P. (2016). ETE 3: Reconstruction, analysis, and visualization of phylogenomic Data. Molecular Biology and Evolution, 33(6), 1635–1638. doi:10.1093/molbev/msw046

51. Hughes, J. F., & Page, D. C. (2015). The biology and evolution of mammalian Y chromosomes. Annual Review of Genetics, 49, 507–527. doi:10.1146/annurev-genet-112414-055311

52. Huson, D. H., Klopper, T., Lockhart, P. J., & Steel, M. A. (2005). Reconstruction of reticulate networks from gene trees. *Research in Computational Molecular Biology*, Proceedings, 3500, 233–249. Retrieved from <Go to ISI>://WOS:000229741100018

53. Jeffries, D. L., Lavanchy, G., Sermier, R., Sredl, M. J., Miura, I., Borzee, A., … Perrin, N. (2018). A rapid rate of sex-chromosome turnover and non-random transitions in true frogs. Nature Communications, 9(1), 4088. doi:10.1038/s41467-018-06517-2

54. Kasjaniuk, M., Grabowska-Joachimiak, A., & Joachimiak, A. J. (2019). Testing the translocation hypothesis and Haldane’s rule in *Rumex hastatulus*. Protoplasma, 256(1), 237–247. doi:10.1007/s00709-018-1295-0

55. Kitano, J., Ross, J. A., Mori, S., Kume, M., Jones, F. C., Chan, Y. F., … Peichel, C. L. (2009). A role for a neo-sex chromosome in stickleback speciation. Nature, 461(7267), 1079–1083. doi:10.1038/nature08441

56. Koenemann, D. M., Kistler, L., & Burke, J. M. (2023). A plastome phylogeny of Rumex (Polygonaceae) illuminates the divergent evolutionary histories of docks and sorrels. Molecular Phylogenetics and Evolution, 182, 107755. doi:10.1016/j.ympev.2023.107755

57. Lahn, B. T., & Page, D. C. (1999). Four evolutionary strata on the human X chromosome. Science, 286(5441), 964–967. doi:10.1126/science.286.5441.964

58. Lenormand, T., Fyon, F., Sun, E., & Roze, D. (2020). Sex chromosome degeneration by regulatory evolution. Current Biology, 30, 3001–3006. doi:10.1016/j.cub.2020.05.052

59. Lenormand, T., Roze, D. (2022). Y recombination arrest and degeneration in the absence of sexual dimorphism. Science, 375(6581), 663–666. doi: 10.1126/science.abj1813

60. Li, H., Durbin, R. (2009). Fast and accurate short read alignment with Burrows-Wheeler transform. Bioinformatics 25(14), 1754–1760. doi: 10.1093/bioinformatics/btp324

61. Li, Z., McKibben, M. T. W., Finch, G. S., Blischak, P. D., Sutherland, B. L., Barker, M. S. (2021). Patterns and processes of diploidization in land plants. Annual Review of Plant Biology, 72, 387–410. doi:10.1146/annurev-arplant-050718-100344

62. Love, A. (1940). Polyploidy in *Rumex acetosella L*. Nature, 145, 351–351. doi:10.1038/145351a0

63. Love, A. (1942). Cytogenetic studies in *Rumex* III. Some notes on the Scandinavian species of the genus. doi: 10.1111/j.1601-5223.1942.tb03281.x

64. Love, A., & Kapoor, B. M. (1967). A chromosome atlas of the collective genus *Rumex*. Cytologia, 32(3-4), 328–342. Retrieved from <Go to ISI>://WOS:A1967C534400004

65. Lovell, J. T., Sreedasyam, A., Schranz, M. E., Wilson, M., Carlson, J. W., Harkess, A., Emms, D., Goodstein, D. M, Schmutz, J. (2022). GENESPACE tracks regions of interest and gene copy number variation across multiple genomes. eLife 11:e78526. doi:10.7554/eLife.78526

66. Lynch, M., & Conery, J. S. (2000). The evolutionary fate and consequences of duplicate genes. Science, 290(5494), 1151–1155. doi:10.1126/science.290.5494.1151

67. Mable, B. K. (2004). ‘Why polyploidy is rarer in animals than in plants’: myths and mechanisms. Biological Journal of the Linnean Society, 82(4), 453–466. doi:10.1111/j.1095-8312.2004.00332.x

68. Maddison, W. P. (1997). Gene trees in species trees. Systematic Biology, 46(3), 523–536. doi: 10.2307/2413694

69. Mendes, F. K., & Hahn, M. W. (2016). Gene tree discordance causes apparent substitution rate variation. Systematic Biology, 65(4), 711–721. doi: 10.1093/sysbio/syw018

70. Mendes, F. K., & Hahn, M. W. (2018). Why concatenation fails near the anomaly zone. Systematic Biology, 67(1), 158–169. doi:10.1093/sysbio/syx063

71. Ming, R., Bendahmane, A., & Renner, S. S. (2011). Sex chromosomes in land plants. Annual Review of Plant Biology, 62, 485–514. doi:10.1146/annurev-arplant-042110-103914

72. Minh, B. Q., Hahn, M. W., & Lanfear, R. (2020). New methods to calculate concordance factors for phylogenomic datasets. Molecular Biology and Evolution, 37(9), 2727–2733. doi:10.1093/molbev/msaa106

73. Minh, B. Q., Schmidt, H. A., Chernomor, O., Schrempf, D., Woodhams, M. D., von Haeseler, A., & Lanfear, R. (2020). IQ-TREE 2: new models and efficient methods for phylogenetic inference in the genomic era. Molecular Biology and Evolution, 37(5), 1530–1534. doi:10.1093/molbev/msaa015

74. Mirarab, S., Reaz, R., Bayzid, M. S., Zimmermann, T., Swenson, M. S., & Warnow, T. (2014). ASTRAL: genome-scale coalescent-based species tree estimation. Bioinformatics, 30(17), i541–548. doi:10.1093/bioinformatics/btu462

75. Mo, Y. K., Lanfear, R., Hahn, M. W., & Minh, B. Q. (2023). Updated site concordance factors minimize effects of homoplasy and taxon sampling. Bioinformatics, 39(1). doi:10.1093/bioinformatics/btac741

76. Muller, J. (1981). Fossil pollen records of extant angiosperms. Botanical Review, 47(1), 1–142. doi:10.1007/Bf02860537

77. Navajas-Perez, R., de la Herran, R., Lopez Gonzalez, G., Jamilena, M., Lozano, R., Ruiz Rejon, C., … Garrido-Ramos, M. A. (2005). The evolution of reproductive systems and sex-determining mechanisms within *Rumex* (polygonaceae) inferred from nuclear and chloroplastidial sequence data. Molecular Biology and Evolution, 22(9), 1929–1939. doi:10.1093/molbev/msi186

78. Ottenburghs, J. (2020). Ghost introgression: spooky gene flow in the distant past. BioEssays, 42(6), e2000012. doi:10.1002/bies.202000012

79. Pala, I., Naurin, S., Stervander, M., Hasselquist, D., Bensch, S., & Hansson, B. (2012). Evidence of a neo-sex chromosome in birds. Heredity, 108(3), 264–272. doi:10.1038/hdy.2011.70

80. Parker, J. S., Wilby, A. S. (1989). Extreme chromosomal heterogeneity in a small-island population of *Rumex acetosa*. Heredity, 62, 133–140. doi:10.1038/hdy.1989.18

81. Pertea, G., & Pertea, M. (2020). GFF Utilities: GffRead and GffCompare. F1000Res, 9. doi:10.12688/f1000research.23297.2

82. Pokorna, M., & Kratochvil, L. (2009). Phylogeny of sex-determining mechanisms in squamate reptiles: are sex chromosomes an evolutionary trap? Zoological Journal of the Linnean Society, 156(1), 168–183. doi:10.1111/j.1096-3642.2008.00481.x

83. Rice, W. R. (1987). The accumulation of sexually antagonistic genes as a selective agent promoting the evolution of reduced recombination between primitive sex chromosomes. Evolution, 41(4), 911–914. doi:10.1111/j.1558-5646.1987.tb05864.x

84. Rifkin, J. L., Beaudry, F. E. G., Humphries, Z., Choudhury, B. I., Barrett, S. C. H., & Wright, S. I. (2021). Widespread recombination suppression facilitates plant sex chromosome evolution. Molecular Biology and Evolution, 38(3), 1018–1030. doi:10.1093/molbev/msaa271

85. Rifkin, J. L., Hnatovska, S., Yuan, M., Sacchi, B. M., Choudhury, B. I., Gong, Y., … Wright, S. I. (2022). Recombination landscape dimorphism and sex chromosome evolution in the dioecious plant *Rumex hastatulus*. Philosophical Transactions of the Royal Society of London B: Biological Sciences, 377(1850), 20210226. doi:10.1098/rstb.2021.0226

86. Sacchi, B., Humphries, Z., Kružlicová, J., Bodláková, M., Pyne, C., Choudhury, B., … Wright, S. I. (2023). Phased assembly of neo-sex chromosomes reveals extensive Y degeneration and rapid genome evolution in Rumex hastatulus. BioRxiv. doi:10.1101/2023.09.26.559509

87. Sanderson, M. J. (2002). Estimating absolute rates of molecular evolution and divergence times: a penalized likelihood approach. Molecular Biology and Evolution, 19(1), 101–109. doi:10.1093/oxfordjournals.molbev.a003974

88. Scrucca, L., Fop, M., Murphy, T. B., & Raftery, A. E. (2016). mclust 5: clustering, classification and density estimation using gaussian finite mixture models. R Journal, 8(1), 289–317. Retrieved from https://www.ncbi.nlm.nih.gov/pubmed/27818791

89. Smith, B. W. (1964). The evolving karyotype of *Rumex hastatulus*. Evolution, 18(1), 93–104. doi:10.2307/2406423

90. Smith, B. W. (1968). Cytogeography and cytotaxonomic relationships of *Rumex paucifolius*. American Journal of Botany, 55(6), 673–683.

91. Smith, M. L., & Hahn, M. W. (2021). New approaches for inferring phylogenies in the presence of paralogs. Trends in Genetics, 37(2), 174–187. doi:10.1016/j.tig.2020.08.012

92. Spigler, R. B., & Ashman, T. L. (2012). Gynodioecy to dioecy: are we there yet? Annals of Botany, 109(3), 531–543. doi:10.1093/aob/mcr170

93. Talavera, M., Balao, F., Casimiro-Soriguer, R., Ortiz, M. A., Terrab, A., Arista, M., … Talavera, S. (2011). Molecular phylogeny and systematics of the highly polymorphic *Rumex bucephalophorus* complex (Polygonaceae). Molecular Phylogenetics and Evolution, 61(3), 659–670.

94. Than, C., Ruths, D., & Nakhleh, L. (2008). PhyloNet: a software package for analyzing and reconstructing reticulate evolutionary relationships. BMC Bioinformatics, 9, 322. doi:10.1186/1471-2105-9-322

95. Tree of Sex Consortium. (2014). Tree of Sex: a database of sexual systems. Scientific Data, 1, 140015. doi:10.1038/sdata.2014.15

96. Tricou, T., Tannier, E., & de Vienne, D. M. (2022a). Ghost lineages can invalidate or even reverse findings regarding gene flow. PLoS Biology, 20(9). doi:10.1371/journal.pbio.3001776

97. Tricou, T., Tannier, E., & de Vienne, D. M. (2022b). Ghost lineages highly influence the interpretation of introgression tests. Systematic Biology, 71(5), 1147–1158. doi:10.1093/sysbio/syac011

98. USDA Agricultural Research Service (2015). Germplasm Resources Information Network (GRIN). USDA Agricultural Research Service. doi:10.15482/USDA.ADC/1212393.

99. Vanderpool, D., Minh, B. Q., Lanfear, R., Hughes, D., Murali, S., Harris, R. A., … Hahn, M. W. (2020). Primate phylogenomics uncovers multiple rapid radiations and ancient interspecific introgression. PLoS Biology, 18(12), e3000954. doi:10.1371/journal.pbio.3000954

100. Vicoso, B. (2019). Molecular and evolutionary dynamics of animal sex-chromosome turnover. Nature Ecology and Evolution, 3(12), 1632–1641. doi:10.1038/s41559-019-1050-8

101. Vicoso, B., & Bachtrog, D. (2013). Reversal of an ancient sex chromosome to an autosome in Drosophila. Nature, 499(7458), 332–335. doi:10.1038/nature12235

102. Wang, Y., Tang, H., Debarry, J. D., Tan., X., Li, J., Wang, X., … Paterson, A. H. (2012). MCScanX: a toolkit for detection and evolutionary analysis of gene synteny and collinearity. Nucleic Acids Research 40(7), e49. doi:10.1093/nar/gkr1293

103. Wernersson, R., & Pedersen, A. G. (2003). RevTrans: Multiple alignment of coding DNA from aligned amino acid sequences. Nucleic Acids Research, 31(13), 3537–3539. doi:10.1093/nar/gkg609

104. White, M. J. D. (1940). The origin and evolution of multiple sex-chromosome mechanisms. Journal of Genetics, 40(1/2), 303–336. doi:10.1007/Bf02982496

105. Yang, Z. (2007). PAML 4: phylogenetic analysis by maximum likelihood. Molecular Biology and Evolution, 24(8), 1586–1591. doi:10.1093/molbev/msm088

106. Yu, Y., & Nakhleh, L. (2015). A maximum pseudo-likelihood approach for phylogenetic networks. BMC Genomics, 16 Suppl 10(Suppl 10), S10. doi:10.1186/1471-2164-16-S10-S10

107. Zhang, C., Rabiee, M., Sayyari, E., & Mirarab, S. (2018). ASTRAL-III: polynomial time species tree reconstruction from partially resolved gene trees. BMC Bioinformatics, 19(Suppl 6), 153. doi:10.1186/s12859-018-2129-y

108. Zhang, L., Li, X., Ma, B., Gao, Q., Du, H., Han, Y., … Qiao, Z. (2017). The tartary buckwheat genome provides insights into rutin biosynthesis and abiotic stress tolerance. Molecular Plant, 10(9), 1224–1237. doi:10.1016/j.molp.2017.08.013

109. Zhou, C., McCarthy, S. A., Durbin, R. (2023). YaHS: yet another Hi-C scaffolding tool. Bioinformatics, 39(1), btac808. doi:10.1093/bioinformatics/btac808

